# Dishevelled-mediated clustering stabilizes Frizzled6 and Vangl2 to establish planar cell polarity in the mammalian skin

**DOI:** 10.64898/2026.05.01.722170

**Authors:** Parijat Sil, Brandon Trejo, Katherine A. Little, Danelle Devenport

## Abstract

Planar cell polarity (PCP) in epithelia is characterized by the polarized distribution of two opposing, membrane-associated PCP complexes across cell junctions. Transmembrane components of the PCP complex bridge cell junctions and organize into punctate, intercellular assemblies that exhibit a high degree of stability. Here, we define the contributions of the cytoplasmic PCP protein, Dishevelled (Dvl), in the sub-micron scale organization and stability of PCP complexes. Using endogenously-tagged fluorescent PCP reporters in the embryonic mouse epidermis, we quantify PCP protein mobility and clustering during polarization. We find that as transmembrane proteins immobilize into puncta, Dishevelled (Dvl2/3) co-accumulates with its transmembrane partner Frizzled (Fz6) in a polarized manner and stabilizes clusters of PCP complexes. We identify a previously unknown function for the oligomerizing DIX domain of Dvl3, typically associated with Wnt signaling, in Dvl3 asymmetric localization. These observations underscore a role for Dvl oligomerization in assembly and stabilization of asymmetric PCP puncta.

## Introduction

Planar cell polarity (PCP) is a fundamental tissue property in which cellular structures and behaviors are aligned along a tissue plane(*1–3*). PCP guides the polarized morphogenesis of tissues and organs and is controlled by a conserved set of membrane-associated protein complexes that localize asymmetrically at cell junctions(*4*). Deciphering how PCP complexes are reorganized from initially uniform to asymmetric localizations is key to understanding how tissues establish planar polarity. Cell biological studies in Drosophila have illustrated how PCP proteins are transported and stabilized over the course of polarization(*4*). To what extent these mechanisms are conserved between insects and mammals is important to resolve.

Studies in the mammalian epidermis have shown that the organization of transmembrane components of the core PCP complex is highly conserved(*5–7*). Similar to wing epithelial cells in Drosophila, the atypical cadherin Fmi/Celsr1 engages in homotypic interactions between cells and partners with Frizzled 6 (Fz6) or Vangl2 on opposing sides of the junction(*4, 5, 7, 8*). The cytoplasmic PCP components, by contrast, are understudied in the context of mammalian planar polarity. In flies, Dishevelled (Dsh) and Diego localize on the Frizzled side of cell junctions whereas Prickle (Pk) localizes with Vang. Dsh and Pk contribute to PCP complex asymmetry, in part, through clustering and/or stabilization of transmembrane complexes(*9–15*). The functions of these cytoplasmic proteins in epidermal planar polarity in mammals have not been explored. For example, there are three Dishevelled (Dvl in vertebrates) homologs in the mammalian genome and it is not known which of these functions in canonical Wnt/β-catenin signaling versus planar polarity establishment in skin.

Dvl proteins associate with Fz receptors in the context of both canonical Wnt signaling and planar polarity, and consist of a DIX, PDZ and DEP domain(*16–18*). The PDZ and DEP domains are required for Fz binding and membrane association, and mutations affecting the DEP domain have been shown to cause PCP-specific defects in Dsh/Dvl function in flies and Xenopus(*19–23*). Oligomerization mediated by the DIX domain assembles Wnt pathway components into membrane-associated ‘Wnt signalosomes’ required for efficient Wnt signal transduction. DIX oligomerization can also generate cytoplasmic condensates that override the requirement for external Wnt ligand to activate signaling(*24–26*). However, the function of the DIX domain in PCP is less clear. Whereas mutations in the DIX domain have little effect on PCP function during convergent extension movements in Xenopus gastrulation and mouse neurulation(*23, 27, 28*), expression of an oligomerization-defective DIX mutant in Drosophila Dsh abrogates Fz and Vang clustering and causes wing hair misalignment(*15*). How Dvl and its oligomerizing DIX domain contribute to the spatial organization of PCP complexes is poorly understood.

Here, we use the mouse epidermis as a mammalian model system to investigate the function of Dvl proteins in the spatial organization of PCP complexes during polarization. Using tissue-specific knockouts, we confirm that mammalian Dvl2 and Dvl3 function as core PCP components in the mouse skin. Combined depletion of Dvl2 and Dvl3 from the skin epithelium results in a loss of Fz6, Vangl2 and Celsr1 asymmetry at epidermal cell junctions and a failure of hair follicles to polarize and align across the anterior-posterior axis. Using FRAP and clustering analyses of endogenously-tagged PCP reporters *in vivo*, we show that loss of polarity in tissues lacking Dvl2/3 function is accompanied by increased turnover and decreased clustering of Fz6 and Vangl2 along epithelial junctions. Further, we show Dvl polarizes concomitant with that of the transmembrane PCP proteins, and accumulates into puncta along cell junctions that are co-enriched with Fz6. *In vivo* structure-function analyses confirm the critical role of the DEP domain in Dvl membrane-recruitment, and show the oligomerizing DIX domain enhances Dvl clustering, puncta formation and asymmetric localization. Our results identify a previously under-appreciated role for Dvl’s oligomerization domain in PCP establishment, offering critical insights into how Dishevelled immobilizes transmembrane PCP components to promote their asymmetric localization.

## Results

### Dvl2 and Dvl3 are required for planar polarity establishment in the murine epidermis

The collective alignment of body hairs across the skin surface arises from the polarized morphogenesis of embryonic hair follicles and is controlled by the core PCP components Celsr1, Fz6 and Vangl2(*5, 29–32*). To determine which of the three mouse Dvl genes is required for epidermal planar polarization, we focused on Dvl2 and Dvl3 because their loss-of-function phenotypes in the inner ear, neural tube, and node are consistent with roles in PCP rather than canonical Wnt signaling(*16, 28, 33, 34*). Additionally, previously published transcriptomic analyses indicate that Dvl2 and Dvl3 are the two predominantly expressed Dvl genes in the skin(*35, 36*). In embryos lacking Dvl3 or Dvl2 individually, via whole-body Dvl3 knockout (*Dvl3−/−*) or epidermal-specific knockout of Dvl2 (*K14-Cre; Dvl2 fl/fl*), hair follicle polarization and PCP protein asymmetry developed normally (Supplementary Figure 1A-D). Combined depletion of Dvl2 and Dvl3 (*K14-Cre; Dvl2 fl/fl; Dvl3−/−*), however, resulted in a severe and complete loss of epidermal planar polarity. Whereas developing hair follicles of wild type embryos polarize along the anterior-posterior (A-P) axis with P-cadherin- and Sox9-expressing cells positioned toward the anterior and posterior, respectively, hair follicles of Dvl2/3 deficient embryos are radially symmetric, with Sox-9 expressing cells forming a ring around central P-cadherin cells (Figure1A-B). Although Dishevelled is a critical component of the Wnt/β-catenin signaling pathway, combined loss of two of the three Dvls did not perturb hair follicle specification or spacing (Figure 1C), suggesting Wnt/β-catenin signaling is unaffected by the loss of Dvl2 and Dvl3 in the epithelial compartment of the skin.

**Figure 1.**
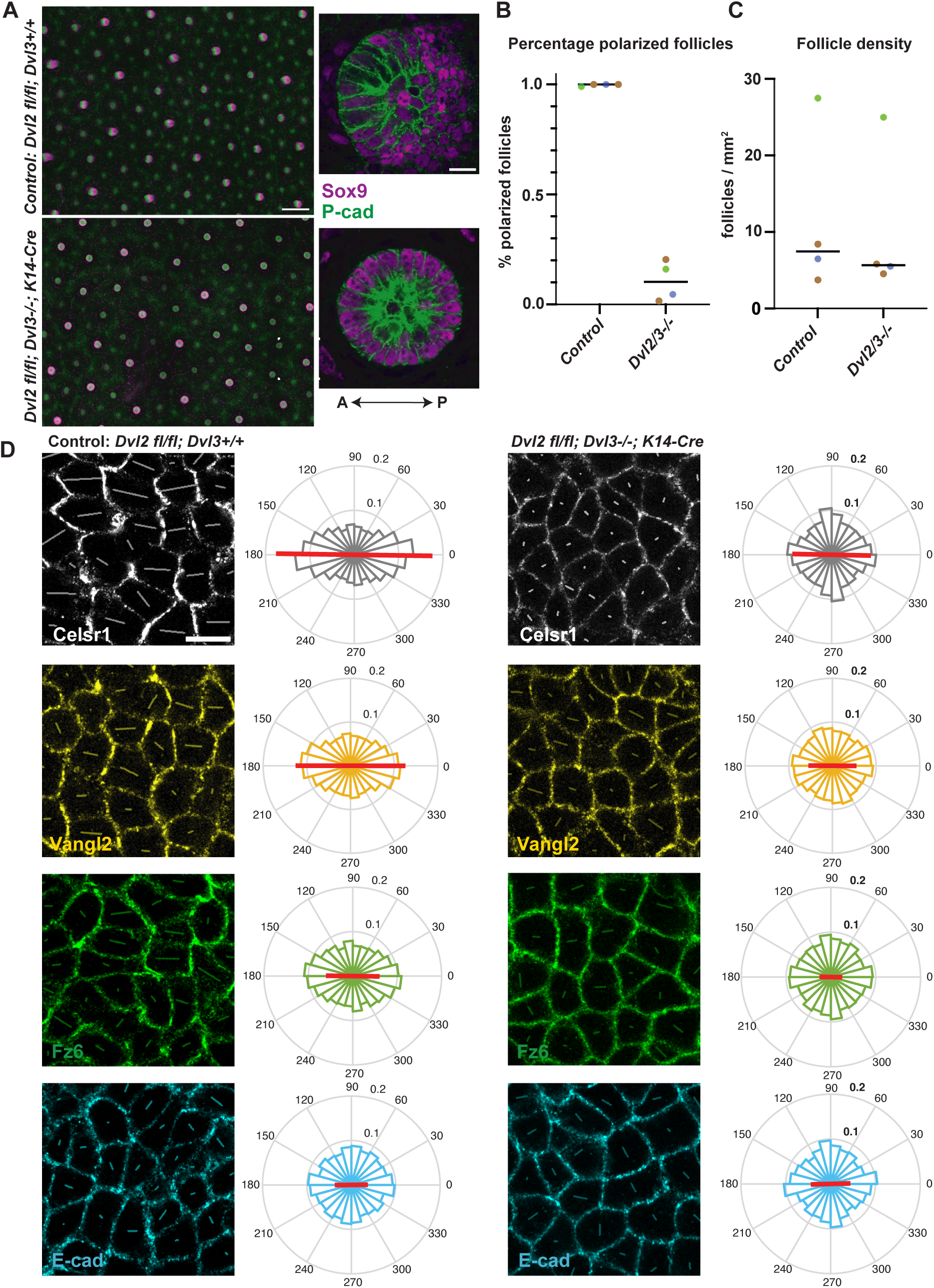
Loss of hair follicle polarization and core PCP protein asymmetry in *Dvl2/3−/−*embryonic skin. **(A)** Average intensity Z-stack projection of E15.5 wholemount backskins from control littermate (*Dvl2fl/fl; Dvl3+/+)*and *Dvl2/3* double mutant *(K14-Cre; Dvl2fl/fl; Dvl3−/−*) embryos immunostained for P-cadherin and Sox9 to label anterior and posterior cell types of developing hair follicles, respectively. Scale bar: 200μm. Zoomed in view of individual follicles imaged at higher magnification shown to the right. Scale bar: 10μm. Anterior is to the left. **(B)** Plot depicting percentage of polarized follicles in littermate controls and *Dvl2/3−/−* double mutant skin (littermates color-matched). **(C)** Plot depicting density of germ-stage follicles in littermate controls and *Dvl2/3*−/− double mutant skin (littermates color-matched). **(D)** Images show planar view of the basal layer of interfollicular epidermis, stained for Celsr1 (gray), Fz6 (green), Vangl2 (yellow) and E-cadherin (cyan). Images are overlaid with polarity nematics indicating the orientation (angle of the line) and magnitude (length of the line) of asymmetry. On the right, circular histograms depicting the angular distribution of polarity. (A-C) 722 follicles from n=4 control littermate embryos and 690 follicles from n=4 *Dvl2/3−/−* embryos. (D) Circular histograms constructed from: n= 7916 basal cells from 4 control embryos and n= 5498 basal cells from 4 *Dvl2/3−/−* embryos stained for Celsr1; n= 9364 basal cells from 4 control embryos and n=10,684 basal cells from 4 *Dvl2/3−/−* embryos stained for E-cadherin; n= 8968 basal cells from 4 control embryos and n= 9456 basal cells from 4 *Dvl2/3−/−* embryos stained for Fz6; n= 9586 basal cells from 4 control embryos and n=10360 basal cells from 4 *Dvl2/3−/−* embryos stained for Vangl2. Scale bar: 10μm

In addition to polarized hair follicle morphogenesis, a hallmark of epidermal planar polarity is the asymmetric localization of the core transmembrane PCP proteins in basal cells of the interfollicular epidermis (IFE). In wild type embryos, Celsr1, Fz6 and Vangl2 are preferentially enriched at anterior-posterior cell junctions and depleted from mediolateral junctions of the dorsal skin, however, combined depletion of Dvl2 and Dvl3 resulted in a complete loss of their polarized distribution. Rather, Celsr1, Fz6 and Vangl2 were uniformly localized along cell junctions of the IFE in *Dvl2; Dvl3* double mutants (Figure 1D). Notably, the epidermal phenotypes displayed in *Dvl2; Dvl3* double mutant embryos are similar to the most severe loss-of-function PCP phenotypes observed in the skin, equivalent to *Celsr1* knockouts(*37*) or *Vangl1; Vangl2* double mutants(*30, 31*). Together, these results demonstrate that Dvl2 and Dvl3 are necessary for epidermal planar polarity in mouse embryos, and that they function redundantly in PCP establishment.

### Dishevelled is asymmetrically localized in the embryonic epidermis

Next, we determined the subcellular localization of Dvl protein in the epidermis over the course of planar polarity establishment. Using an antibody that recognizes both Dvl2 and Dvl3, we found the distribution of Dvl2/3 in basal cells of the IFE became increasingly polarized within the epithelial plane from E13.5 to E15.5 stages of development (Figure 2A-B). At E13.5, Dvl2/3 was mostly uniformly distributed around the lateral edges of basal cells, and became more concentrated to A-P edges at E15.5 (Figure 2A). The timing and progression of Dvl2/3 asymmetry mirrored that which we have previously reported for Celsr1, Fz6 and Vangl2(*5, 38*). However, we noted that compared to the transmembrane components, Dvl2/3 displayed a more cytoplasmic distribution in basal cells, especially at earlier developmental stages. To quantify this, we measured the colocalization between Dvl2/3 and Fz6, as well as E-cadherin, an epidermal protein that is membrane-associated but non-planar polarized. Colocalization between Dvl2/3 and Fz6 increased between E13.5 and E15.5 whereas colocalization between Dvl2/3 and E-Cadherin did not, indicating that Dvl2/3 becomes increasingly membrane-associated likely by specific interactions with Fz6. Moreover, colocalization between Dvl2/3 and Fz6 was greater than colocalization with E-cadherin at both early (E13.5) and late (E15.5) stages, reflecting a basal Dvl2/3 and Fz6 co-enrichment which is enhanced upon polarity establishment (Figure 2C, Supplementary Figure 2C-D).

**Figure 2.**
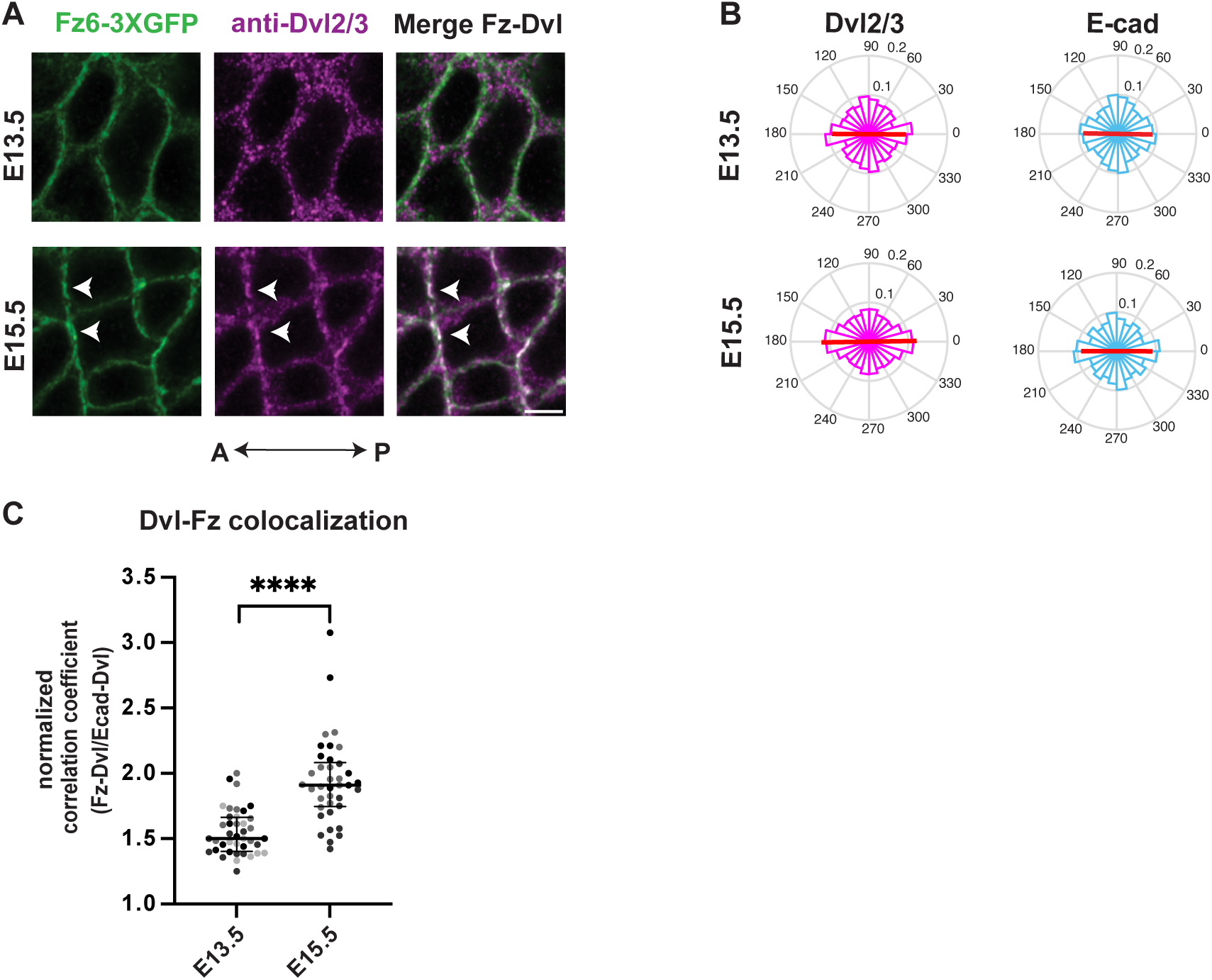
Dvl2/3 becomes asymmetrically localized at epidermal cell junctions over developmental time. **(A)** Planar view of the epidermal basal layer of E13.5 (top panel) and E15.5 (bottom panel) wholemount skin immunostained for Fz6 (green) and Dvl2/3 (magenta) and merged as indicated. White arrowheads point to regions of colocalization between Dvl2/3 and Fz6. Scale bar: 5μm. **(B)** Circular histograms of angular distribution of Dvl2/3 asymmetry show polarization is higher at E15.5 compared to E13.5, while polarization of E-cadherin is unchanged. n= 5604 basal cells from 3 embryos at E13.5 and n= 3480 basal cells from 2 embryos at E15.5 stained for E-cadherin; n= 6220 basal cells from 3 embryos at E13.5 and n= 5581 basal cells from 2 embryos at E15.5 stained for Dvl2/3. **(C)** Plot of normalized correlation coefficient with median and interquartile range. Pearson’s correlation coefficient of Dvl2/3-Fz6 is normalized to that of Dvl2/3-E-cadherin (as proxy for extent of Dvl membrane association). n= 40 ROIs from 4 embryos at E13.5 and n= 38 ROIs from 3 embryos at E15.5. **** indicates p<0.0001 by KS test

We also examined the localization of Dvl1, and found the antibody labeled only a subset of cells within the center of developing hair follicles (Supplementary Figure 2A-B). The absence of Dvl1 from the IFE, together with the localization and function of Dvl2/3 in the skin is consistent with the conclusion that Dvl2 and Dvl3 are the primary Dvl genes involved in epidermal planar polarity establishment.

### Dishevelled is required for Fz6 and Vangl2 immobilization within punctate assemblies

To investigate how Dvl2/3 might contribute to the asymmetric localization of the transmembrane PCP components, we began by characterizing features of Fz6 and Vangl2 organization and dynamics over the course of planar polarity establishment. Fluorescence recovery after photobleaching (FRAP) was used to measure Fz6 and Vangl2 turnover and mobility along cell junctions at unpolarized (E13.5) and polarized (E15.5) stages (Figure 3 and Figure S3). FRAP analysis was performed in epidermal explants from embryos expressing endogenously-tagged Fz6-3XGFP and tdT-Vangl2(*6*). At E13.5, both proteins were uniformly distributed along A-P and mediolateral (M-L) junctions and displayed large mobile fractions at both types of junctions (Figure S3A-D). By contrast, at E15.5, Fz6-3XGFP and tdT-Vangl2 were non-uniformly distributed along A-P edges, forming puncta with higher fluorescence intensity interspersed by low intensity regions (Figure 3A-B). Within puncta, Fz6-3XGFP and tdT-Vangl2 were largely immobile (∼60% or higher immobile fraction for both proteins) whereas in non-puncta regions, their mobile fractions were similar to those at E13.5 A-P junctions as well as M-L junctions at E15.5 (Figure 3G and 3J). To assess whether the inherent speed of diffusion of the protein is different between M-L and A-P junctions or within and outside the puncta regions, the half-life of recovery was measured. Only the half-life of Fz6-3XGFP within puncta, but not tdT-Vangl2, was modestly higher compared to non-puncta regions of A-P junctions (Figure S3E-F). Thus, concomitant with their asymmetric localization, Fz6 and Vangl2 become immobilized selectively at A-P junctions, accumulating as a stable pool of proteins as they incorporate into puncta.

**Figure 3.**
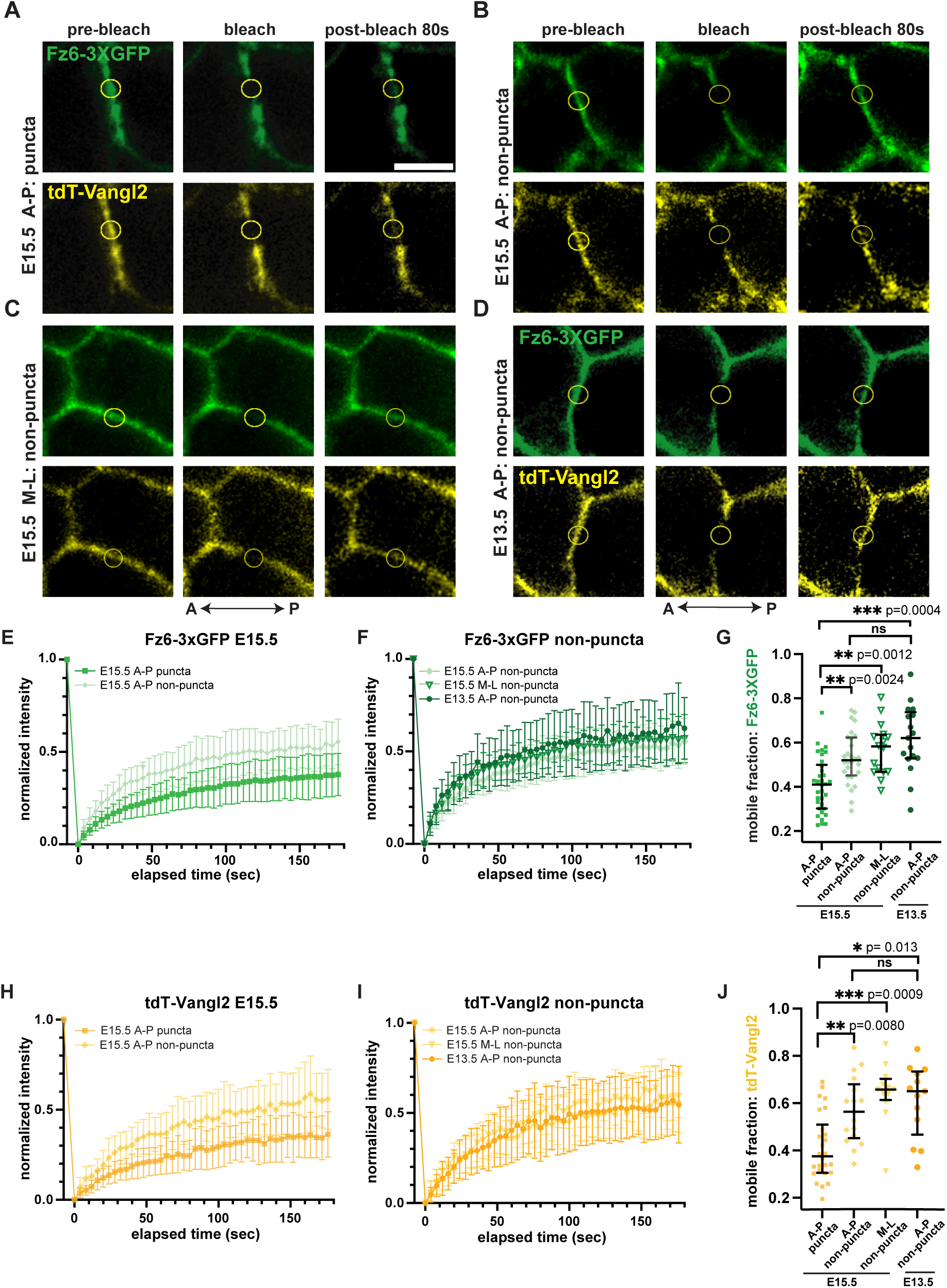
Fz6 and Vangl2 are immobilized within puncta along A-P junctions during polarity establishment. **(A-D)** Still images from FRAP time course performed on skin explants from *Fz6-3XGFP; tdTomato-Vangl2* embryos at the indicated stages. Fz6-3XGFP (green) and tdT-Vangl2 (yellow) are shown at pre-bleach, bleach (0 sec) and post-bleach (80 sec) time points. Yellow circles mark bleached regions analyzed for fluorescence recovery. FRAP analysis was performed on (A) punctate region of an A-P junction in E15.5 epidermis, (B) non-puncta region of an A-P junction in E15.5 epidermis, (C) non-puncta region of an mediolateral (M-L) junction in E15.5 epidermis, and (D) non-puncta region of an A-P junction in E13.5 epidermis. **(E and H)** FRAP recovery curves of Fz6-3XGFP (E) and td-Tomato Vangl2 (H) at puncta and non-puncta regions of A-P junctions in E15.5 epidermis. Mean and standard deviation are shown. Note the higher mobile fraction of Fz6 and Vangl2 in non-puncta regions. **(F and I)** FRAP recovery curves of Fz6-3XGFP (F) and tdTomato-Vangl2 (I) from non-puncta regions of A-P and M-L junctions in E15.5 epidermis and non-puncta regions of A-P junctions in E13.5 epidermis. Mean and standard deviation are shown. Note similar mobile fractions within non-puncta regions irrespective of junction orientation and developmental stage. n= 36 puncta for Fz6 and 32 puncta for Vangl2 at A-P junctions at E15.5. n= 32 non-puncta A-P traces at E15.5, n=20 non-puncta regions of M-L junctions at E15.5 and n= 22 non-puncta A-P traces at E13.5 for each of Fz6-3XGFP and tdTomato-Vangl2. **(G and J)** Mobile fractions of Fz6-3XGFP in E and F (G) and tdTomato-Vangl2 in H and I (J). Median and inter-quartile range are shown. Mobile fractions were computed only from the subset of traces that fitted a one-phase association equation with a chi squared value of ≥ 0.8. Scale bar: 5μm.

To test whether Dvl is required for stabilization of Fz6 and Vangl2, Fz6-3XGFP and tdT-Vangl2 reporters were crossed in *Dvl2/3* double mutants. In E15.5 epidermal explants from *Dvl2/3* double mutants, Fz6-3XGFP and tdT-Vangl2 (*K14-Cre; Dvl2 fl/fl; Dvl3 −/−; Fz6-3XGFP; tdT-Vangl2)* were no longer concentrated into high intensity puncta along A-P junctions (Figure 4E). Furthermore, their mobile fractions calculated from FRAP measurements were significantly higher than controls and more similar to unpolarized junctions of wild-type embryos at E13.5 and of non-puncta regions of A-P junctions of embryos at E15.5 (Figure 4E-H, S4E-F). Consistently, the recovery half-lives of non-puncta regions of control junctions and *Dvl2/3* double mutant junctions were similar (Figures S4G-H).

**Figure 4.**
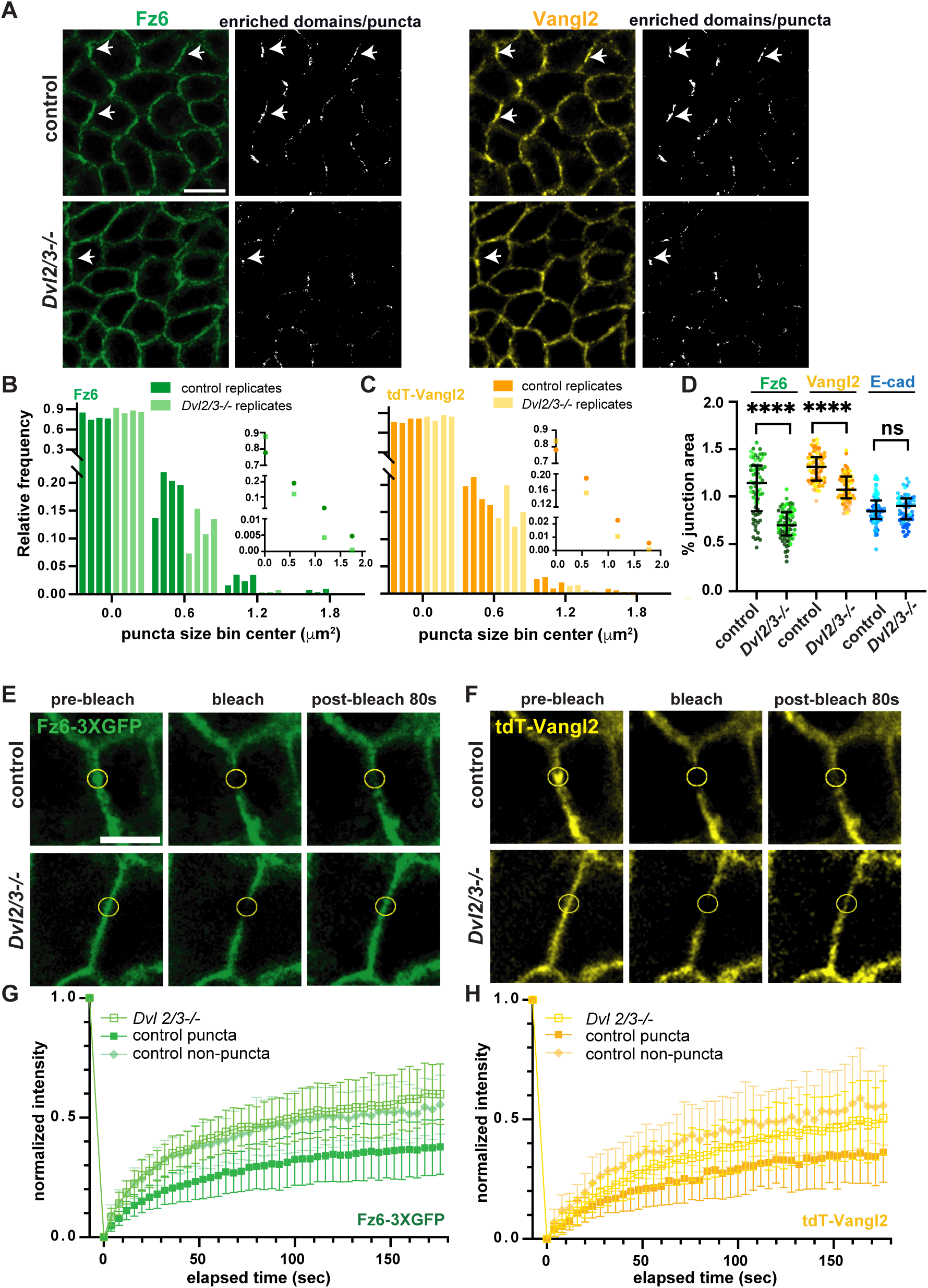
Clustering of Fz6 and Vangl2 along junctions requires Dvl2 and Dvl3. **(A)** Planar views of basal layer of E15.5 embryonic skin immunostained for Fz6 (green) and Vangl2 (yellow). Right panels show corresponding segmented images depicting spatially enriched domains/puncta of Fz6 and Vangl2. **(B,C)** Normalized frequency distribution of Fz6 (B) and Vangl2 (C) puncta sizes along epidermal junctions. Bars represent frequencies of individual control or *Dvl2/3−/−* replicates, n=4 embryos. Insets show normalized frequency distribution of puncta sizes pooled from 4 control and 4 *Dvl2/3−/−* epidermis. p<0.0001 by Chi-Square test. n= 2546 Fz6 puncta and n=2853 Vangl2 puncta pooled from 4 control embryos, n=2383 Fz6 and n= 2932 Vangl2 puncta pooled from 4 *Dvl2/3−/−* embryos. **(D)** Percentage of junctional area occupied by puncta of Fz6, Vangl2 and E-cad, a non-PCP control transmembrane protein, in control versus *Dvl2/3−/−* epidermis. n= 69 ROIs pooled from 4 control embryos and n= 71 ROIs pooled from 4 *Dvl2/3−/−* embryos. **** indicates p<0.0001 by KS test. **(E, F)** Still images from FRAP time course of Fz6-3XGFP (E) and tdTomato-Vangl2 (F) at pre-bleach, bleach (0 sec) and post-bleach (80 sec) time points. Yellow circles mark photobleached regions analyzed for fluorescence recovery. A-P junctions of E15.5 control and *Dvl2/3−/−* epidermis were sampled. **(G, H)** FRAP recovery curves of junctional Fz6-3XGFP (G) and tdTomato-Vangl2 (H) at puncta and non-puncta regions of control and non-puncta regions *Dvl2/3 −/−* epidermis. n= 36 Fz6 puncta and 32 Vangl2 puncta in control epidermis pooled from 4 embryos, n= 32 non-puncta A-P traces pooled from 3 control embryos, and n= 56 non-puncta regions of A-P junctions pooled from 4 *Dvl2/3−/−* embryos for each of Fz6 and Vangl2. Scale bars: 10μm in A and 5μm in E and F.

The enrichment of core PCP components into puncta along cell junctions is a conserved feature of PCP organization in epithelial tissues and has been observed in both live and fixed samples and by super resolution imaging(*6–8, 10, 15, 39*). To characterize the non-uniform organization of Fz6 and Vangl2 in the epidermis, we immunolabeled E15.5 wild-type epidermal explants to simultaneously visualize Fz6 and Vangl2 at high resolution, and measured puncta size, fractional area of all junctions covered by Fz6 and Vangl2 puncta, and degree of enrichment of Fz6 and Vangl2 within puncta. In wild-type epidermis, the majority of Fz6 and Vangl2-containing puncta ranged in size from ∼ 0.1 to 1um^2^ (Figures 4A-C, Figure S4C) and typically covered <1.5% of epidermal junctional area (Figure 4D). In contrast, puncta sizes were significantly smaller in epidermal tissues lacking Dvl2/3 and these smaller puncta were distributed uniformly around both A-P and M-L junctions (Figures 4A-C, Figure S4C). Specifically, puncta in the size range of ∼0.5 um^2^ and larger were significantly reduced in the mutant compared to the control epidermis (Figures 4B-C). These puncta showed lower enrichment of Fz6 and Vangl2 and covered a smaller fraction of the epidermal junctional area (Figures 4D and S4D). E-cadherin puncta, in contrast, did not exhibit a reduction in size or junctional area coverage in the absence of Dvl2/3, indicating that the effect of Dvl2/3 loss on clustering is selective for PCP components (Figures-4D, S4A-D). The change in Fz6 and Vangl2 punctate organization likely arises from a failure to become immobilized in the absence of Dvl2/3 function (Figures 4E-4H). Together, these mobility and puncta analyses conclusively demonstrate that a key function of Dvl in PCP establishment is to stabilize assemblies of transmembrane components into puncta.

### The DEP and DIX domains are required for Dishevelled recruitment to and enrichment at cell junctions

To begin to decipher the functions of Dvl’s structural domains, we performed a structure-functional analysis of mouse Dvl3 in cultured keratinocytes. Primary mouse keratinocytes were transfected with either full length Dvl3, or a Dvl3 variant lacking either its DEP domain, DIX domain, or C-terminal intrinsically disordered (IDR) domain (Figure 5A-5B). Each variant was co-transfected with Celsr1 and Fz6, which are both required for Dvl3 recruitment to keratinocyte junctions. Whereas full length Dvl3 was recruited to Fz6-enriched interfaces in which two Celsr1 expressing cells had engaged in homotypic adhesion, Dvl3βDEP failed to enrich at Celsr1-Fz6 interfaces and remained diffusely localized in the cytoplasm. Removal of the C-terminal IDR domain did not have an effect on junctional recruitment, but deletion of the DIX domain reduced Dvl3 enrichment (Figure 5C). Fz6 localization to junctions was comparable across all conditions, indicating that the differences in Dvl recruitment observed was not due to differential levels of Fz6 junctional enrichment (Figure S5A).

**Figure 5.**
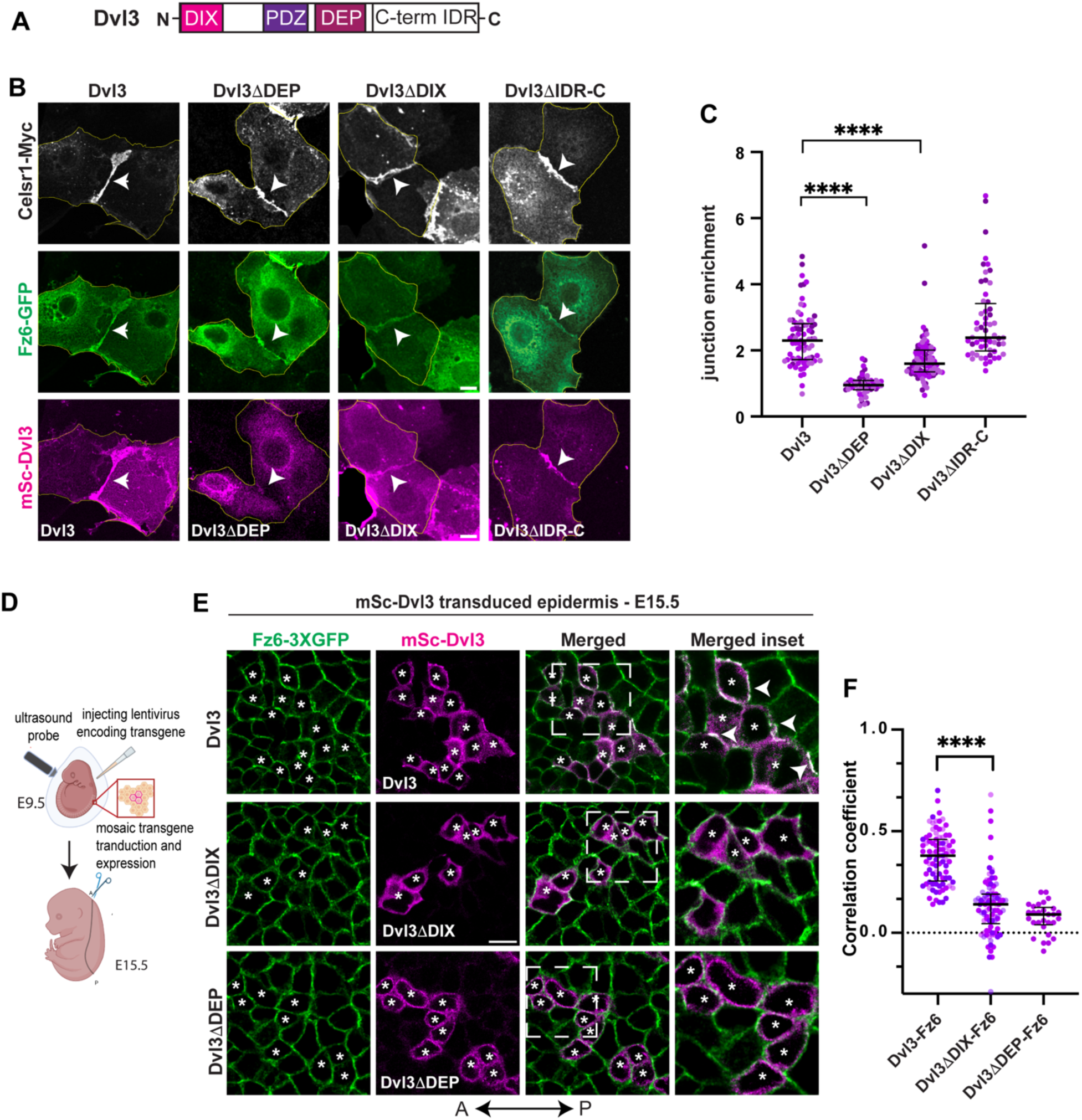
DEP and DIX domain interactions are required for Dvl junctional localization. **(A)** Schematic of Dvl3 protein domains. (B) Representative examples of keratinocytes transfected with Celsr1-Myc, Fz6-GFP and either full-length mScarlet3-Dvl3 (mSc-Dvl3) or deletion mutants Dvl3ΔDEP, Dvl3ΔDIX, Dvl3ΔIDR-C. **(C)** Junction enrichment ratios with median and inter-quartile range of full length and deletion variants of Dvl3. n= 75 junctions for Dvl3, 50 junctions for Dvl3ΔDEP, 88 junctions for Dvl3ΔDIX and 56 junctions of Dvl3ΔIDR-C, pooled from three replicates. **(D)** Schematic of the workflow of intra-uterine lentiviral transduction of E9.5 embryos to transduce the surface ectoderm for visualization of mSc-Dvl3 variants at E15.5. **(E)** Representative examples of epidermal basal layer from e15.5 embryos transduced with lentivirus carrying either full-length mSc-Dvl3, mSc-Dvl3ΔDEP, or mSc-Dvl3ΔDIX (magenta) and co-stained for Fz6 (green). White asterisks mark transduced cells in the basal layer. Merged insets with arrowheads indicate colocalization of mSc-Dvl3 with Fz6. **(F)** Pearson’s correlation coefficients between Fz6 and either mSc-Dvl3, mSc-Dvl3ΔDEP, or mSc-Dvl3ΔDIX as a measure of colocalization. Median and interquartile range are shown. n=88 cell clusters from 3 transduced embryos for mSc-Dvl3, n=93 cell clusters from 4 transduced embryos for mSc-Dvl3ΔDIX and n=29 cell clusters from 1 transduced embryo for mSc-Dvl3ΔDEP. ****: p<0.0001, tested by KS-test. scale bar: 10μm.

To confirm these data *in vivo*, we transduced the embryonic epidermis with lentivirus carrying mScarlet-tagged full-length Dvl3, Dvl3βDEP or Dvl3βDIX deletion variants using ultrasound-guided *in utero* lentiviral injections(*40*) (Figure 5D). Following injection at E9.5, embryos were harvested at E15.5 for whole mount imaging. Similar to our observations in cultured keratinocytes, Dvl3βDEP completely failed to localize to the membrane and showed poor colocalization with endogenous Fz6. Dvl3βDIX, by contrast, was membrane-localized but displayed significantly lower co-enrichment with Fz6 compared to full length Dvl3 (Figure 5E-5F). Together, these results suggest that the DEP and DIX domains of Dvl3 perform distinct functions in planar polarization. The DEP domain is critical for membrane and/or Fz6 binding whereas the oligomerizing DIX domain enhances junctional localization.

### The DIX domain is required for Dvl3 clustering and asymmetry

Thus far we have shown that Dvl2/3 are required to cluster and stabilize core PCP components into asymmetrically localized puncta. Given the known ability for DIX domains to oligomerize into large multimers, we asked whether Dvl3 lacking its DIX domain might be defective in its ability to become asymmetrically localized. Similar to the behavior of Fz6 and Vangl2 in the epidermis from E13.5 to E15.5, Dvl2/3 becomes increasingly clustered and asymmetrically enriched into puncta at A-P junctions (Figures 6A-6D). To test the role of the DIX domain in its spatial reorganization, we measured the asymmetry and puncta formation of mScarlet-Dvl3 expressed mosaically in lentiviral transduced epidermis. In mosaic patches of transduced Dvl3-expressing cells, Dvl3 was enriched on the posterior edges of single cells or cell clusters, where Fz6 is known to reside (Figures 6H-6I), and was organized into puncta (Figures 6D-6G). In contrast, Dvl3βDIX did not cluster to the same extent as full length Dvl3 (Figures 6D-6G), nor was it localized preferentially to the posterior side of cell edges (Figures 6H-6I).

**Figure 6.**
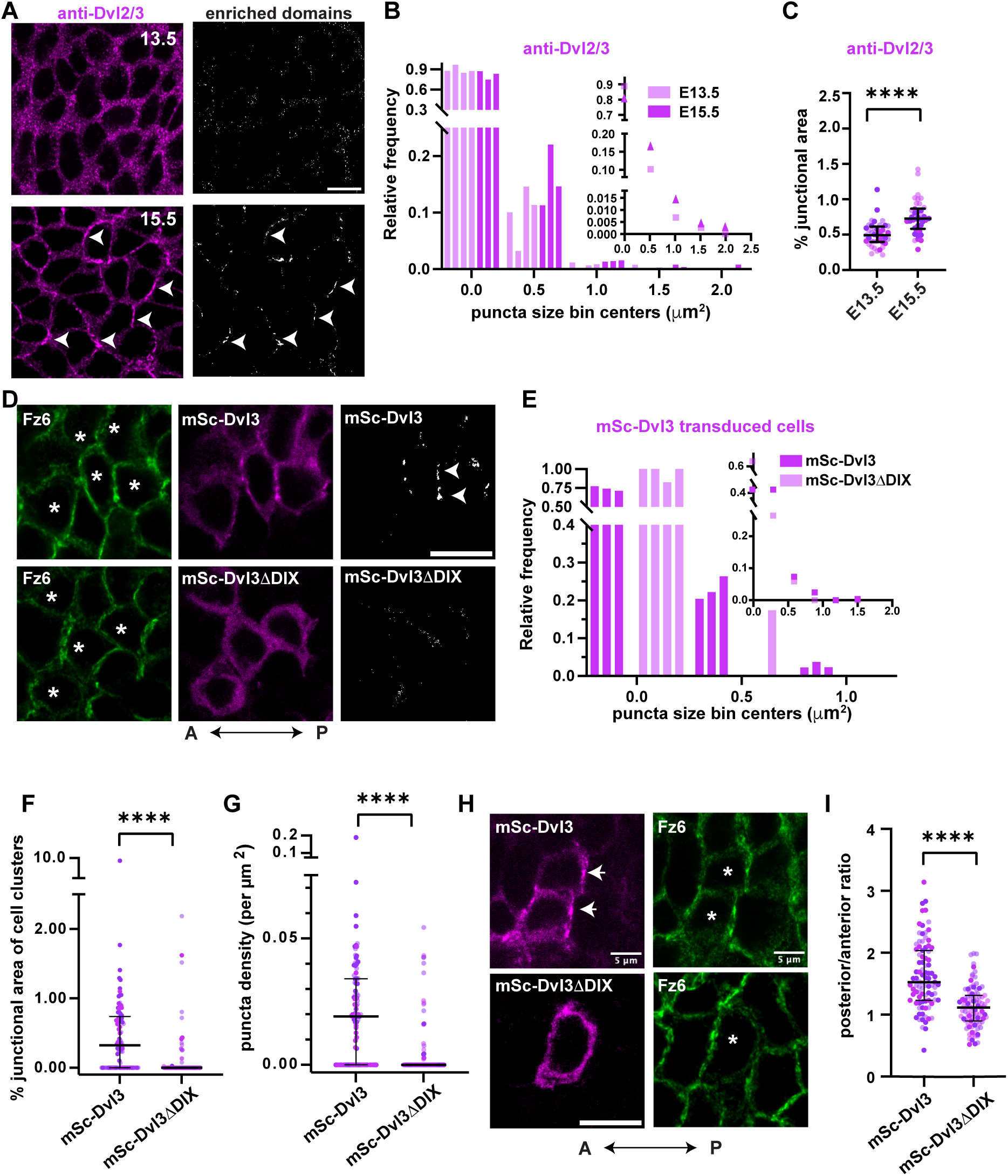
Polarized accumulation of Dvl3 requires interactions mediated by the DIX domain. **(A)** Representative planar views of the epidermal basal layer at E13.5 and E15.5 labeled with anti-Dvl2/3 antibody (magenta). Right panels show segmented areas of Dvl2/3 accumulation into enriched domains/puncta. Arrowheads point to puncta of Dvl2/3 at E15.5. **(B)** Normalized frequency distribution of Dvl2/3 puncta sizes with each bar representing one replicate of E13.5 (light pink) and E15.5 (dark pink) epidermis from 4 and 3 embryos, respectively. The inset shows normalized frequency distribution of pooled puncta sizes. p<0.0001 by Chi-Square test. n= 1159 puncta/enriched domains from 4 embryos at E13.5 and 1304 puncta/enriched domains from 3 embryos at E15.5. **(C)** Percent junctional area covered by Dvl2/3 puncta/enriched domains at E13.5 and E15.5. Median and inter-quartile range are shown. n= 40 ROIs from 4 embryos at E13.5 and n=38 ROIs from 3 embryos at E15.5. **(D)** Representative planar view of E15.5 epidermal basal layer transduced with mSc-Dvl3 or mSc-Dvl3ΔDIX (magenta), and immunostained for Fz6 (green). Right panels and arrowheads show segmented puncta/enriched domains. **(E)** Quantification of normalized frequency distribution of mSc-Dvl3 and mSc-Dvl3ΔDIX puncta sizes. Note only mSc-Dvl3 forms large puncta. Each bar represents one replicate of 3 embryos transduced with mSc-Dvl3 (dark pink) and 4 embryos transduced with mSc-Dvl3ΔDIX. Inset shows normalized frequency distribution of puncta sizes pooled from 3 and 4 embryos transduced with mSc-Dvl3 and mSc3-Dvl3ΔDIX, respectively. n= 245 puncta from 89 transduced cell clusters or single cells across 3 embryos for mSc3-Dvl3; n= 34 puncta from 93 cell clusters or single cells across 4 embryos for mSc3-Dvl3ΔDIX. **(F)** Percent junctional area occupied by puncta/enriched domains of mSc-Dvl3 or mSc-Dvl3ΔDIX, with median and interquartile range. p<0.0001, KS test. **(G)** Puncta density (# detected puncta per unit junctional area) for mSc-Dvl3 and mSc-Dvl3ΔDIX, with median and interquartile range. p<0.0001, KS test. **(H)** Images of basal cells transduced with mSc-Dvl3 or mSc-Dvl3ΔDIX highlighting polarized accumulation of mSc-Dvl3 along posterior edges of cell clusters (white arrowheads). Transduced cells marked by asterisks in Fz6 channel. **(I)** Fluorescence intensity ratio of mSc-Dvl3 and mSc-Dvl3ΔDIX at posterior versus anterior junctions with median and inter-quartile range. n=91 cells for mSc-Dvl3 from 3 embryos and n=95 cells for mSc-Dvl3ΔDIX from 4 embryos. p<0.0001 by KS test. Scale bars: 10μm.

Owing to the low probability of transducing double *Dvl2/3* knockout embryos to conduct rescue experiments, we were unable to test the ability of DIX domain mutants to rescue Dvl2/3 loss of function. Nevertheless, these results confirm a role for the oligomerizing DIX domain in Dvl’s asymmetric localization and suggests Dvl oligomerization may play a central role in the stabilizing interactions leading to planar polarization of core PCP components.

## Discussion

The assembly of protein complexes at cell surfaces underlies the formation of specialized membrane compartments that function in cell adhesion, signal transduction, and cell polarity(*41–46*). How cytoplasmic scaffolding proteins interact with and organize their transmembrane partners into higher order assemblies is key to understanding polarized membrane organization. Dishevelled is an essential scaffolding protein in the Wnt/β-catenin pathway that organizes Wnt pathway components into multiprotein, membrane-associated signalosomes(*16, 24, 47*). Whether Dvl serves a similar function in the organization of planar cell polarity pathway components is less understood. Moreover, the contribution of Dvl family proteins to mammalian PCP establishment were unknown. Here, using *in vivo* protein diffusion assays and protein clustering measurements in mouse embryonic epidermis, we show that Dishevelled is essential for the transition of PCP protein mobility and clustering toward an immobile, punctate state, properties that are associated with their reorganization from a uniform to asymmetric localization.

Despite the known roles for Dvl proteins in both Wnt signaling and planar polarity, our data strongly suggest that Dvl2 and Dvl3 play redundant roles in epidermal planar cell polarity but not Wnt signaling. Wnt signaling through β-catenin-mediated transcription is essential for specifying hair follicle fate(*48, 49*), and we did not observe defects in hair follicle specification, downgrowth or spacing in embryos lacking epidermal Dvl2/3 function. We conclude that Dvl1 can mediate canonical Wnt signaling functions in the epidermis, at least at embryonic stages examined here.

The asymmetric accumulation of membrane-associated polarity proteins involves the movement of proteins from initially uniform distributions and the retention of proteins at asymmetric positions. Through FRAP mobility measurements we have shown that both Fz6 and Vangl2 freely diffuse within the membrane at E13.5 and later at E15.5 are trapped in punctate assemblies when asymmetry is established. Half-life recovery measurements of Fz6 and Vangl2, which serve as a proxy for the diffusion coefficient, were similar between non-punctate regions at E15.5 and unpolarized junctions at E13.5 suggesting the mobile protein pool does not undergo significant transitions in their diffusion behavior across the different developmental stages. A lack of difference in half-life of recovery in puncta and non-puncta regions of A-P junctions but a significant increase in immobile pool within puncta-containing regions likely suggest that PCP proteins can associate to build a puncta in a manner where they are not readily exchanged with the surroundings in the measured timescale. These observations mirror aspects of junctional PCP protein dynamics measured by FRAP in Drosophila(*10, 39*) and are consistent with a mechanism in which PCP proteins cluster via a filament-type assembly process, as recently proposed from single-molecule counting experiments (*15*). Finally, the dramatic loss of immobilization and accumulation of Fz6 and Vangl2 in puncta in the *Dvl2/3* double knockout epidermis is consistent with observations made in Drosophila wing epithelia(*11*, *29*, *38*), pointing to a conserved role for Dsh/Dvl proteins in polarity establishment across diverse epithelial tissues and species (*10, 15, 39*).

Early structure-function analyses of Dsh/Dvl in both flies and vertebrates lead to the prevailing view that the DEP domain performs critical PCP functions whereas the DIX domain predominantly functions in Wnt signaling responses(*20–23, 50*). For example, in mouse neurulation, the DIX domain is dispensable for its membrane localization and PCP function(*28*). Similarly, early studies in flies suggested only a minor role for the DIX domain in wing hair polarization(*22*). More recently, however, point mutations in the DIX domain of Drosophila Dsh were shown to impair Fz and Vang assembly at junctions as well as their segregation to opposite sides of wing epithelial cells(*14, 15*). Additionally, recent evidence revealed that Dact1-dependent Dvl2 oligomerization via the DIX domain is critical for convergent extension in Xenopus embryos, wherein Dact1 promotes co-accumulation of Dvl2 and Fz7 along cell boundaries in animal cap explants(*51*). Through the expression of Dvl3 deletion variants in the developing embryonic skin, we establish a role for the oligomerizing DIX domain in Dvl3’s polarized accumulation. Oligomerization can serve to enhance the avidity of otherwise weak protein-protein interactions, increasing binding dwell times and slowing protein turnover(*25, 47*). Given the slow turnover of PCP proteins residing in puncta, it is tempting to speculate that Dsh/Dvl utilizes DIX-dependent oligomerization to concentrate and stabilize Fz-Dvl complexes in puncta, which, in a non-cell autonomous manner, stabilizes Vangl accumulation in neighbors. In this manner, DIX-based oligomerization may play a conserved function in building polarized membrane domains in the context of PCP and in the formation of Wnt signalosomes.

## Materials and Methods

### Mouse lines and breeding

Mice were housed according to the guide for the Care and Use of Laboratory Animals in an AAALAC-accredited facility. Animal husbandry and maintenance were carried out in complete compliance with Laboratory Animal Welfare Act. All of the procedures involving mice, including protocol for surgery and post-op care were approved by Princeton University’s Institutional Animal Care and Use Committee (IACUC). The Dvl3 mutant 129S-*Dvl3^tm1Awb^*/J (Strain #:009083)(*33*) and Dvl2flox mouse B6;129-*Dvl2^tm1.1Wds^*/J, (Strain #:029061)(*52*) mice were obtained from Jackson laboratories. Fz6-3XGFP and tdT-Vangl2 mice were at Princeton, and were generated in a previous study(*6*). K14-Cre mouse line originated from the laboratory of Elaine Fuchs(*53*). C57BL/6 mice were used for staging immunofluorescence experiments. All genotyping were carried out with Transnetyx using custom designed probes. Experiments were performed on E13.5 or E15.5 embryos (irrespective of male or female) and obtained by mating heterozygous mice specifically for Dvl3 mutants and tdT-Vangl2 carrying strains.

### Immunofluorescence of embryonic skin

Embryonic back-skins were fixed and stained using published protocols(*37*). Briefly, E15.5 embryos were fixed for 1h at room temperature (RT) in 4% paraformaldehyde in PBS supplemented with CaCl2 and MgCl2. Following three washes in PBS for 30 minutes each, full thickness dorsal skin explants were dissected in PBS. All antibody stainings were performed in a blocking solution containing 1% bovine serum albumin (BSA), 1% fish gelatin, 2% goat serum and 2% donkey serum (for anti-Fz6, 4% donkey serum was used in place of goat serum) in PBS containing 0.2% Triton-X100. PBS was replaced by TBS when staining for P-Cadherin. Following overnight incubation in blocking buffer at 4°C, skins were washed five times with PBS or TBS containing 0.2% Triton and incubated overnight at 4°C with primary antibodies. Samples were washed again five times in PBS or TBS containing 0.2% Triton, followed by 2 hours of incubation with secondary antibody (1:1000) at RT or overnight at 4°C. Samples were finally washed four times with PBS or TBS containing 0.2% Triton, first of which was supplemented with 1ug/ml Hoechst (Cat: H1399) to stain nuclei. Samples were mounted with Prolong Gold. The following primary antibodies were used at the specified dilutions: guinea pig anti-Celsr1 (Danelle Devenport, 1:1000)(*5*), goat anti-Fz6 (1:400, R&D Biosystems, Cat: AF1526), rabbit anti-E-cadherin (1:250, Cell Signaling: 3195), rat anti-Vangl2 (1:100, Millipore, Cat: MABN750), rabbit anti-Dvl3 (Proteintech 13444-1-AP, 1:400), rabbit anti-Sox9 (Millipore, AB5535, 1:1000), rat anti-P-cadherin (Clontech, M109, 1:200). Alexa Fluor −488, −555, and −647 secondary antibodies were used at 1:1000 (Invitrogen or Jackson ImmunoResearch).

### Imaging of embryonic skin and hair follicle polarity analysis

For hair follicle polarity analysis, images were acquired using a Nikon A1R-STED confocal microscope operated by NIS Elements software, using PlanApo 20X 0.75 NA air objective with resonance scanning mode. High resolution hair follicle images were acquired with 60X 1.45 NA oil immersion objective and galvo scanning. 20X images were then stitched in NIS Elements and processed in Fiji. Average Intensity Projection (AIP) images were generated for angle calculations using the angle tool in ImageJ. 60X images were processed in Fiji.

### PCP protein asymmetry imaging and analysis

For analysis of core PCP protein asymmetry in the basal layer, images were acquired on Nikon A1R-Si confocal microscope (for wildtype and mutant analysis) and on Nikon A1R STED confocal microscope (for staging polarization analysis) controlled by NIS Elements software using PlanApo 60 × 1.45 NA oil immersion objective. Cellpose(*54*) was used to segment cell edges of the basal layer. Segmentation masks were obtained for each channel, and masks were post-processed, and hand corrected in ImageJ if required. Polarity analysis was performed using Tissue Analyzer V2 plugin in ImageJ(*55*). Tissue Analyzer used the segmentation masks generated in Cellpose to calculate axis and magnitude (nematic order) of membrane localized proteins (as defined in (*55, 56*)). Circular histograms of the data were plotted in MATLAB. Images were rotated to align them along the anterior-posterior axis before analysis.

### Keratinocyte culture, transfection

Wildtype CD1 or *Fz6−/−* keratinocytes were cultured in E-Media prepared in the laboratory according to published protocol(*57*). Media was supplemented with 50 µM CaCl2. Cells were transfected using Effectene reagent (Qiagen #301425) following a modified manufacturer’s protocol: 300 ng DNA was used to transfect each well of 12-well plates. For co-transfection with three different plasmids, one of which was Celsr1-Myc, a ratio of 2:1:1 of Celsr1: plasmid2: plasmid3 was used. See Table 1 for full list of plasmids used in this study.

**Table 1.** Plasmid details.

| Name | Backbone details | Insert details | Source |
| --- | --- | --- | --- |
| Celsr1-Myc | Stratagene pCMV-3Tag-9 | Full length mouse Celsr1 tagged to Myc tag on the C terminus | (5) |
| Fz6-GFP | K14 MCS vector | Full length mouse Fz6 tagged to GFP on the C terminus | Made by Danelle Devenport in Fuchs laboratory |
| mSc-Dvl3 | Lentiviral, EF1-alpha promoter for mSc-Dvl3 expression. Backbone also | mScarlet3 and 3X-FLAG tag at the N- | Generated for this paper |
|  | expresses Cre under PGK promoter | terminus of Dvl3 |  |
| mSc-ΔDEP | Lentiviral, EF1-alpha promoter for mSc-Dvl3ΔDEP expression. Backbone also expresses Cre under PGK promoter | mScarlet3 and 3X-FLAG tag at the N-terminus of Dvl3. DEP domain deleted | Generated for this paper |
| mSc-Dvl3ΔDIX | Lentiviral, EF1-alpha promoter for mSc-Dvl3ΔDIX expression. Backbone also expresses Cre under PGK promoter | mScarlet3 and 3X-FLAG tag at the N-terminus of Dvl3. DIX domain deleted | Generated for this paper |
| mSc-Dvl3ΔIDR-C | Lentiviral, EF1-alpha promoter for mSc-Dvl3ΔIDR-C expression. Backbone also expresses Cre under PGK promoter | mScarlet3 and 3X-FLAG tag at the N-terminus of Dvl3. C terminal IDR deleted. | Generated for this paper |

### Junction enrichment assay

For junction enrichment assay, ∼100,000 keratinocytes were seeded on fibronectin coated #1.5 glass coverslips in each well of 12-well plate in low calcium E-media (50 µM CaCl2). ∼24 h post-plating, cells were transfected with necessary plasmid combinations. ∼24 h post-transfection, low calcium E-media was replaced with 1.5 mM CaCl2 containing E-media. After ∼24 h of incubation in high calcium media, cells were fixed and stained.

### Immunofluorescence of keratinocytes

Confluent keratinocyte monolayers were rinsed in PBS containing Calcium and Magnesium chloride(PBS++) and fixed with 4% PFA in PBS++ for 10 min at RT, followed by permeabilization in PBS containing 0.1% Triton-X 100 (PBT1) for 10 min at RT. Primary antibodies were diluted 1:1000 for anti-Myc and 1:2000 for anti-GFP in PBT1 and cells were incubated with the same for 30 min. Following primary antibody treatment, cells were washed three times in PBS for five minutes each and further incubated for 30 min with secondary antibodies and Hoechst (1:2000) in PBT1. Cells were washed three times in PBS and mounted on glass slides using Prolong Gold.

### Image acquisition and junction enrichment analysis in cultured keratinocytes

Cells were imaged using a PlanApo 20 × 0.75NA Air objective with additional zoom on a Nikon A1R-STED confocal microscope equipped with 405, 488, 561, and 643 nm lasers. Each channel was sequentially imaged to avoid bleed-through. Maximum intensity projections of the Z stacks were generated in Fiji/ImageJ and background subtracted. ROIs were drawn along the junctions marked by Celsr1-Myc. Another ROI was made along the periphery of the transfected cells sharing the junction. A ratio was obtained of the background corrected mean intensities of the junction and the cell pair/individual cell ROI as follows:

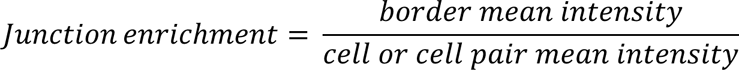

Data were extracted and processed in Microsoft Excel and plotted using GraphPad Prism, were statistical analysis was performed.

### Fluorescence recovery after photobleaching

E13.5 or E15.5 backskins from embryos expressing Fz6-3XGFP and tdT-Vangl2 (in wild type or Dvl2/3−/− backgrounds) were mounted on agar pads supplemented with F-media as previously described before for live imaging of whole-mount back skins. Live skin explants were imaged on a Nikon A1R-STED confocal with a 100X oil immersion objective with 1.45 NA and a pixel size of 70nm was used for acquisition. A 405nm laser was used to photobleach both Fz6-3XGFP and tdT-Vangl2 simultaneously. Magnification, laser power (for bleach and acquisition), pixel dwell time and acquisition rate were kept uniform across all measurements. 1um diameter circular ROIs were created to bleach and sample A-P or M-L junctions in puncta or non-puncta regions as applicable. FRAP acquisition sequence consisted of 3 reference pre-bleach images with no intervals, followed by bleach (4.8s) and 45 frames with 4-s intervals to monitor fluorescence recovery. The acquired images in the time series were checked for Z-drift and corrected for XY-drift and discarded when significant drift was present. Three reference ROIs were made along non-bleached junctions to correct for overall bleaching during image acquisition. The ROI values were extracted using NIS elements software for subsequent analysis in Microsoft Excel and Graphpad Prism. Each image time series was bleach corrected (hereafter referred as corrected intensity) and corrected intensity profile was normalized as *(F t –F bleach)/(F ini – F bleach)*, where, Ft =corrected intensity of the ROI at a given time point, Fbleach = corrected intensity at the time point immediately after bleaching, Fini = mean ROI intensity of three pre-bleach frames. Mean as well as individual recovery curves were fitted to exponential one phase association equation in Graphpad Prism to determine half-life of recovery and the fitted Plateau and Y0 values were used to determine the *mobile fraction* = *(Plateau-Y 0)/(1-Y 0)*. The averaged traces for each condition were fitted to the model with an r-squared value >0.9 and individual traces with r-squared value >0.8 were included for generating mobile fraction and half-life plots.

### Image acquisition and analysis for puncta characterization

For puncta analysis of PCP components, Z-stacks of whole mounted epidermal explants were acquired using 100X, 1.45 NA oil immersion objective on Nikon-A1R confocal STED microscope, with a pixel size of 98 nm and a Z-step size of 0.4 micrometer. Z-projections of three optimal focal planes in which the basal keratinocyte layer was in focus, were used to generate maximum intensity projection images. Images were background corrected with mean intensity measured from the junctions of unstained samples or of un-transduced areas of samples (as applicable) imaged with the same settings. The junction network of a given ROI was segmented using the Mean preset within the Threshold dialog of ImageJ with adjustments to the min and max as needed for reliable detection of junctions. This was done for each fluorescent channel to create a mask of protein localization along all junctions. Mean intensity and standard deviation values for the entire junctional network was extracted and the mean + 3 standard deviations was calculated for each ROI. This value was used as a threshold value and regions in the image with pixel values above this were designated as puncta. Puncta above this threshold intensity that were also equal to or greater than 9 pixel^2^ in area (to disregard any cluster that was not Nyquist-sampled by at least 3 pixels in any direction, assuming a square shape) were analyzed to calculate puncta size, junctional area covered by puncta (total area covered by puncta in an ROI/ junction network area in the ROI) and fold enrichment of protein in puncta (mean intensity of all clusters in an ROI/ junctional mean intensity in the ROI). Data were extracted and processed in Microsoft Excel and plotted using GraphPad Prism, were statistical analysis was performed.

### Lentivirus production and *in utero* lentiviral injections

Lentiviruses were produced in HEK-293T cells using published protocol(*40*). Briefly, HEK cells were seeded for transfection and cultured in D10 media (DMEM (10-017-CV)+10% v/v FBS+ 1% V/V Pen-Strep/L-glut mix+ 1% v/v 7.5% Sodium bicarbonate+ 1% v/v 100mM Sodium Pyruvate+G418) in 500cm^2^ poly-L-lysine coated plates (two plates for each prep). Confluent HEK cell monolayers were co-transfected using packaging plasmids: 275βg psPax2, 180 βg pmd2.g and 275βg of either 3X-FLAG-mScarlet-Dvl3, 3X-FLAG-mScarlet-Dvl3βDEP, or 3X-FLAG-mScarlet-Dvl3βDIX transgene expressing lentiviral plasmid (for plasmid details see table) using calcium chloride in HBSS. Transfection mix was diluted in D10 without G418. 14 hours post transfection, transfection media was replaced with D10 without G418. 2 hours later (16 hours post transfection), media was replaced by Virus production media (Freestyle media^TM^ 293+ 1% v/v pen-strep/ L-glut mix+1% v/v 100mM sodium pyruvate+ 1% v/v 7.5 % sodium bicarbonate and 1% 5mM sodium butyrate). 46 hours post transfection, supernatants were collected and filtered to remove cell debris. Viral supernatants were concentrated using low speed centrifugation at 3300g at 4°C through 100kDa MW cut-off Millipore Centricon 70 Plus (Fisher UFC7 100-08) cartridge to concentrate 140ml of supernatant to <1ml. The concentrated supernatant was then ultracentrifuged over a sucrose cushion at 45000 RPM in Beckman Coulter SW-50.1 rotor for 90 minutes at 4°C and resuspended in ∼60μl to achieve a ∼2000 fold concentration of virus with minimal titer loss.

Lentiviral injections were performed following established ultrasound-guided in utero amniotic injection protocols adapted from Beronja et al(*40*) under approved institutional IACUC guidelines. Pregnant mice at embryonic day 9.5 (E9.5) were anesthetized with isoflurane and maintained under anesthesia for approximately 45 min. Pre-operative analgesia was provided by subcutaneous injection of meloxicam, and Ethiqa XR was administered for extended post-operative pain control. A midline laparotomy was performed under sterile surgical conditions to expose the uterus, and two embryos at a time from a single uterine horn were gently exteriorized into a sterile phosphate-buffered saline (PBS) filled dish (VisualSonics catalog #SA-11620) for visualization. Embryos were visualized by ultrasound using a VisualSonics Vevo 3100 LT imaging system mounted on a dedicated imaging station (VisualSonics P/N #11228-01). Concentrated lentivirus (∼1.2 µl per embryo) was injected into the amniotic cavity using pulled glass capillary needles (FUJI FILM catalog #PIP35-BV20-DNJ) connected to a microinjector system (Drummond nanoject II catalog #3-000-204) under ultrasound guidance. In each surgery, four embryos were injected, while remaining embryos in the two horns were left uninjected and used as matched littermate controls. During injections, uninjected embryos were maintained within the abdominal cavity to minimize exposure. Following injection, uterine horns were returned to the abdominal cavity, the abdominal wall was sutured closed, and the skin incision was secured with surgical staples. Dams were recovered on a heating pad until fully awake and mobile, then monitored daily for five days post-surgery. Meloxicam was administered for two additional days post-operatively, with Ethiqa XR given on post-operative day two. Embryos were harvested at E15.5 for downstream analyses.

### Statistics

Details of sample size, statistical significance, p-values are described in figures and legends of each figure. Individual data points are displayed in all plots to depict spread and different Ns are color coded in the plots wherever applicable. Differences between distributions of junction enrichment ratios, correlation co-efficient, mobile fraction and half-life distribution plots, puncta size and related parameters were tested by non-parametric KS test or Chi-square test as applicable, using Graphpad Prism software. Significance of difference in recovery curves in FRAP were determined from non-overlap of confidence intervals of the mean fitted curves.

## Acknowledgements

We thank Devenport lab members for their discussion and feedback and in particular Rishabh Saran for fruitful discussions on data analysis. We thank Gary Laevsky and Sha Wang of the Confocal Core Facility at Princeton University, a Nikon Center for Excellence, for help with imaging set-up and guidance. We thank Ricardo Mallarino and Sarah Mereby at Princeton University and Carlos Patino Descovich from MSKCC for assistance with in utero lentiviral transduction procedures.

Research reported here was supported by National Institute of Arthritis and Musculoskeletal and Skin Diseases (NIAMS) of the National Institutes of Health under award number R01AR066070 and R01AR068320, the Eunice Kennedy Shriver National Institute of Child Health and Human Development (NICHD) under grant number R01HD105009, and the New Jersey Commission on Cancer Research under grant number: COCR24PDF018, The content is solely the responsibility of the authors and does not necessarily represent the official views of the National Institutes of Health.

## Author contributions

PS conceptualized the study, set up all the assays and performed all the experiments and data analyses. BT set up and performed lentivirus injection procedures on mice. KAL helped with reagent generation for the study. PS and DD secured funding for the project. DD supervised the study by overseeing all aspects of experiment design, data generation, interpretation and analysis. PS and DD wrote and edited the manuscript.

