## Supplementary information for "Dishevelled-mediated clustering stabilizes Frizzled6 and Vangl2 to establish planar cell polarity in the mammalian skin"

### Figure S1

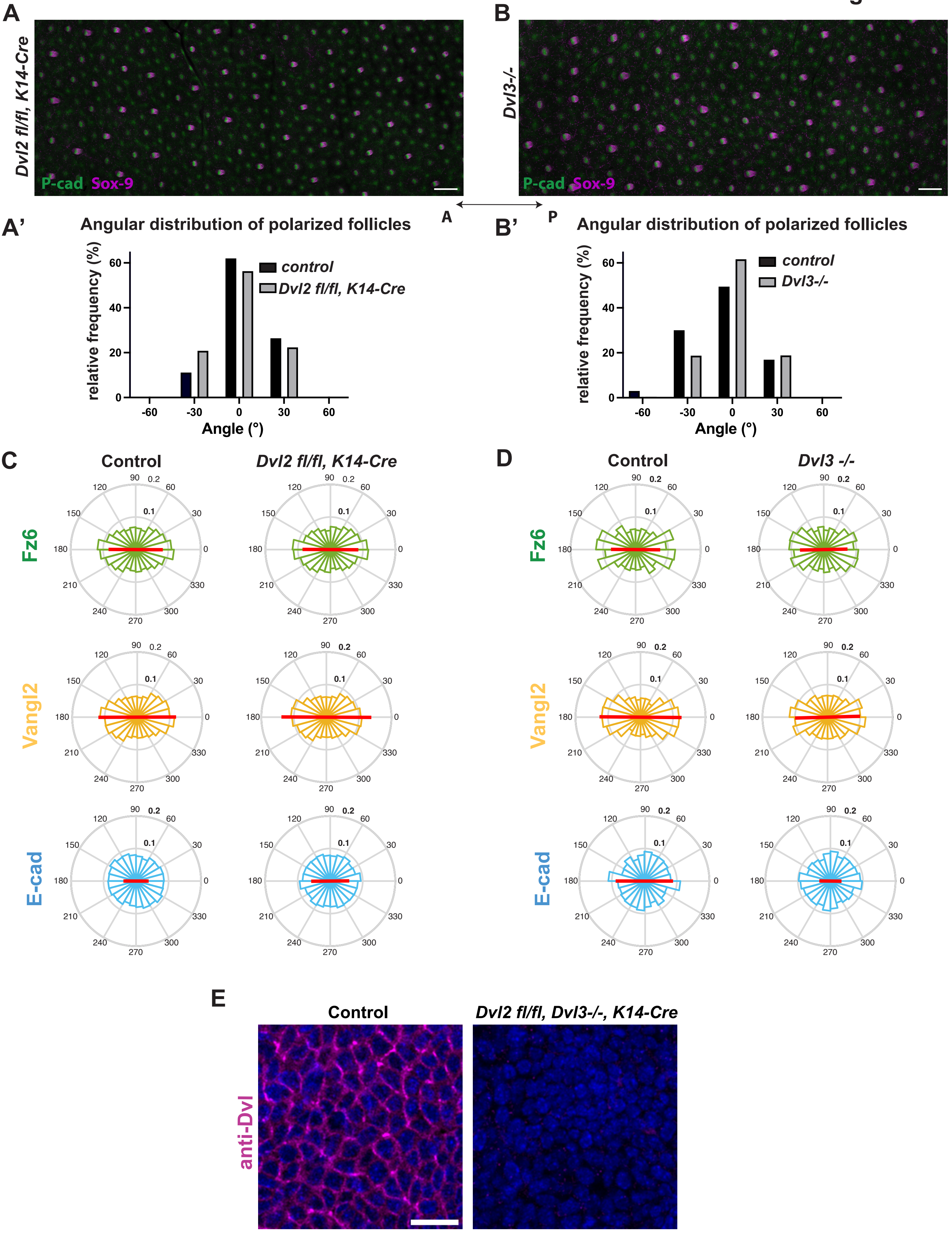
A
Dvl2 fl/fl, K14-Cre
P-cad Sox-9
B
Dvl3-/-
P-cad Sox-9
A
P
A'
Angular distribution of polarized follicles
relative frequency (%)
control
Dvl2 fl/fl, K14-Cre
Angle (°)
B'
Angular distribution of polarized follicles
relative frequency (%)
control
Dvl3-/-
Angle (°)
C
Control
Dvl2 fl/fl, K14-Cre
Fz6
Vangl2
E-cad
D
Control
Dvl3 -/-
Fz6
Vangl2
E-cad
E
Control
Dvl2 fl/fl, Dvl3-/-, K14-Cre
anti-Dvl

**Supplementary Figure 1. Hair follicle polarization and core protein asymmetry is unperturbed in *Dvl2* and *Dvl3* individual knockouts. (A-B)** Average intensity projection of Z stack images of E15.5 backskin lacking Dvl2 (*Dvl2 fl/fl; K14-Cre*, A) or lacking Dvl3 (*Dvl3-/-,* B) immunostained for P-cadherin and Sox9 to label anterior and posterior cells of developing hair follicles, respectively. Scale bar: 200μm. **(A'-B')** Histograms showing angular distribution of polarized hair follicles in *Dvl2fl/fl; K14-Cre* and *Dvl3-/-* epidermis. **(C-D)** Circular histograms depicting the angular distribution of polarity of Fz6, Vangl2 and E-cadherin in *Dvl2fl/fl; K14-Cre* (C) and *Dvl3-/-* epidermis (D) compared to littermate controls. Circular histograms in (C) constructed from: n= 4962 basal cells from 2 control embryos and n=9190 basal cells from 4 *Dvl2fl/fl; K14-Cre* embryos stained for Fz6; n= 4624 basal cells from 2 control embryos and n= 6976 basal cells from 4 *Dvl2fl/fl; K14-Cre* embryos of stained for Vangl2; n= 4170 basal cells from 2 control embryos and n= 8234 basal cells from 4 *Dvl2fl/fl; K14-Cre* embryos stained for E-cadherin. Circular histograms in (D) constructed from: n= 1820 basal cells from 1 control embryo and n= 2944 basal cells from 2 *Dvl3-/-* embryos stained for Fz6; n= 1722 basal cells from 1 littermate control and n= 3404 basal cells from 2 of *Dvl3-/-* embryos stained for Vangl2. n= 1552 basal cells from 1 littermate control embryo and n= 3060 basal cells from 2 *Dvl3-/-* embryos stained for E-cadherin. **(E)** Planar view of the epidermal basal layer of littermate control (left) and *Dvl2fl/fl; K14-Cre; Dvl3-/-* (right) stained with anti-Dvl 2/3 antibody (magenta signal in interfollicular epidermis) shows a loss of Dvl2 and Dvl3 in the double knockout skin. Scale bar: 10μm.

**Figure S2**

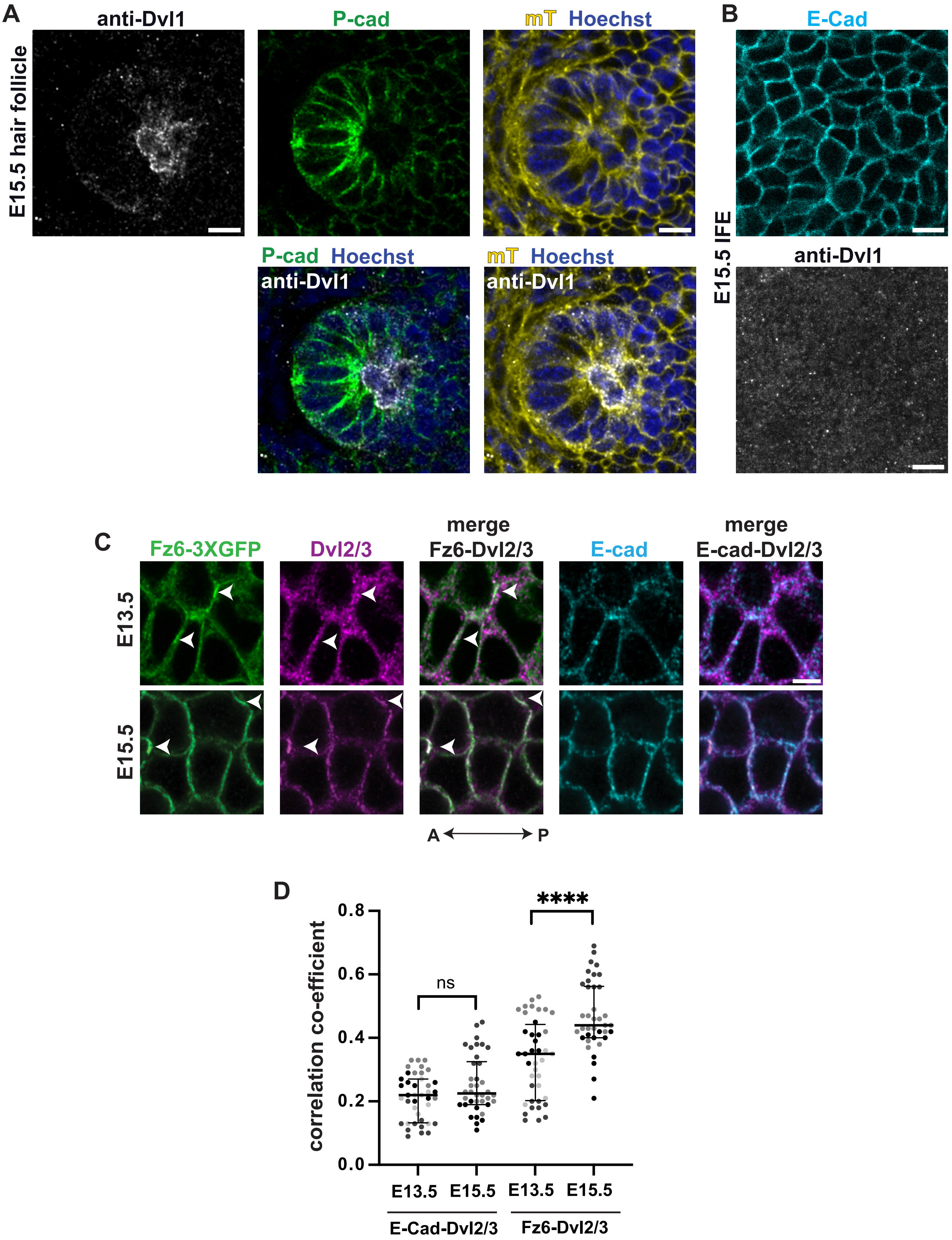
A
anti-Dvl1
P-cad
mT Hoechst
E15.5 hair follicle
P-cad Hoechst
anti-Dvl1
mT Hoechst
anti-Dvl1
B
E-Cad
anti-Dvl1
E15.5 IFE
C
Fz6-3XGFP
Dvl2/3
merge
Fz6-Dvl2/3
E-cad
merge
E-cad-Dvl2/3
E13.5
E15.5
A
P
D
correlation co-efficient
0.8
0.6
0.4
0.2
0.0
ns
****
E13.5
E15.5
E13.5
E15.5
E-Cad-Dvl2/3
Fz6-Dvl2/3

**Supplementary Figure 2. Unlike Dvl1, Dvl2/3 are expressed and localized with Fz6 in the IFE. (A)** Representative image of germ-stage hair follicle from E15.5 epidermis labeled for Dvl1 (grayscale), P-cadherin (green), Hoechst (blue) and membrane-tdTomato (mT; yellow). Note Dvl1 is restricted to cells in the centre of the follicle. Scale bar: 10μm. **(B)** Planar view of the basal epidermal layer at E15.5 labeled with E-cadherin (cyan, top) and Dvl1 (grayscale bottom) antibodies. Note Dvl1 is absent from the IFE marked by E-Cadherin. Scale bar: 10μm **(C)** Planar views of the basal layer of E13.5 (top panel) and E15.5 (bottom panel) whole mount epidermis stained for Fz6 (green), Dvl2/3 (magenta) and E-cadherin (cyan) with merged overlays as indicated. White arrowheads point to regions of Dvl and Fz6 colocalization. Scale bar: 5μm. **(D)** Plot of Pearson's correlation coefficient of Dvl2/3 with Fz6 and Dvl2/3 with E-cadherin, with median and inter-quartile range, n= 40 ROIs from 4 embryos at E13.5 and n= 38 ROIs from 3 embryos at E15.5. **** indicates $p<0.0001$ by KS test.

### Figure S3

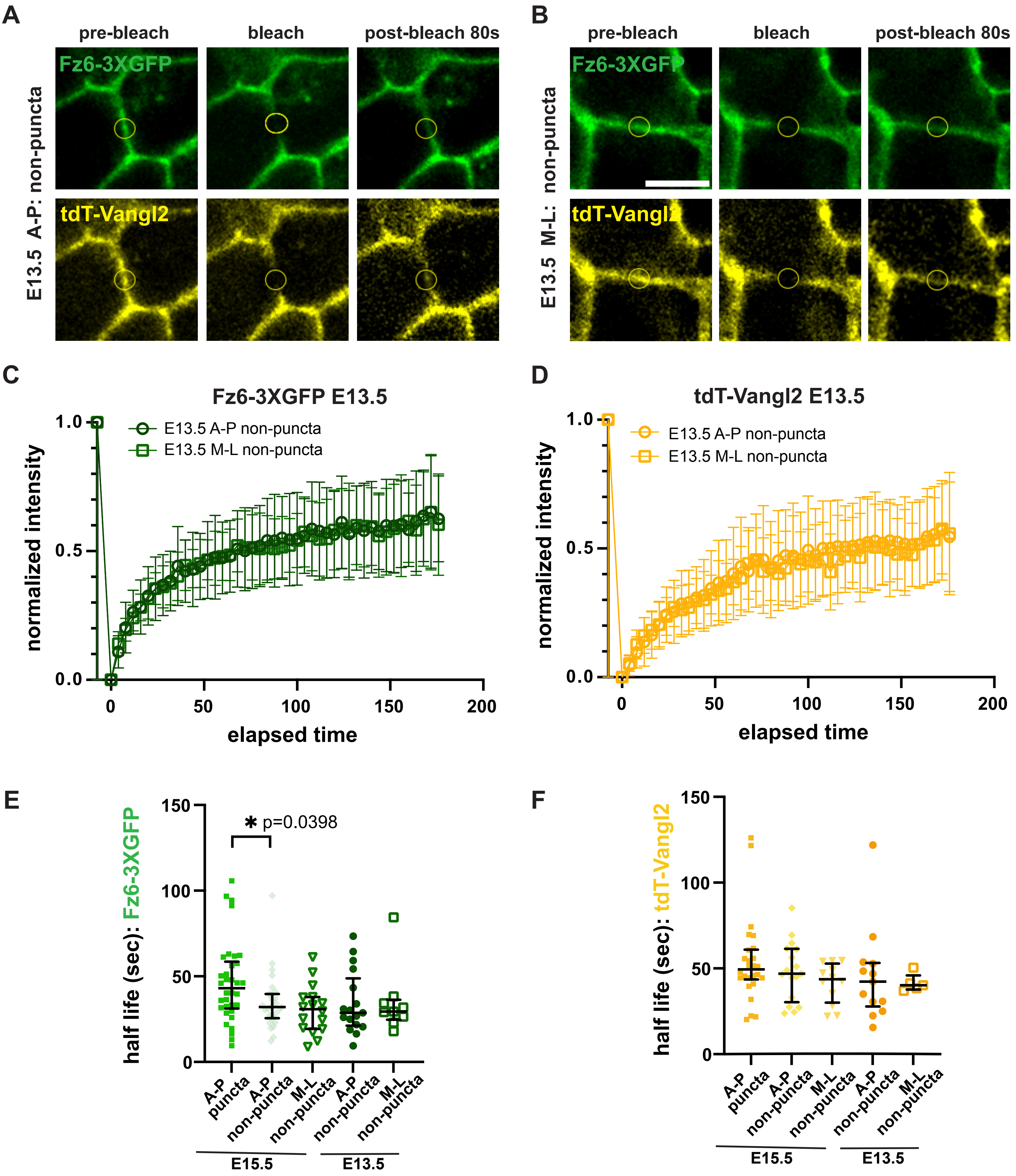
A
pre-bleach
bleach
post-bleach 80s
Fz6-3XGFP
tdT-Vangl2
E13.5 A-P: non-puncta
B
pre-bleach
bleach
post-bleach 80s
Fz6-3XGFP
tdT-Vangl2
E13.5 M-L: non-puncta
C
Fz6-3XGFP E13.5
E13.5 A-P non-puncta
E13.5 M-L non-puncta
normalized intensity
elapsed time
D
tdT-Vangl2 E13.5
E13.5 A-P non-puncta
E13.5 M-L non-puncta
normalized intensity
elapsed time
E
half life (sec): Fz6-3XGFP
* p=0.0398
A-P puncta
A-P non-puncta
M-L non-puncta
A-P non-puncta
M-L non-puncta
E15.5
E13.5
F
half life (sec): tdT-Vangl2
A-P puncta
A-P non-puncta
M-L non-puncta
A-P non-puncta
M-L non-puncta
E15.5
E13.5

**Supplementary Figure 3. FRAP analysis of Fz6-3XGFP and tdT-Vangl2. (A-B)** Still images from FRAP time course of Fz6-3XGFP and tdTomato-Vangl2 at pre-bleach, bleach (0 sec) and post-bleach (80 sec) time points. Yellow circles depict photobleached regions analyzed for fluorescence recovery at (A) non-puncta region of A-P junction in E13.5 epidermis, (B) non-puncta region of M-L junction in E13.5 epidermis (B). **(C, D)** FRAP recovery curves of Fz6-3XGFP (C) and tdTomato-Vangl2 (D) at non-puncta regions of A-P and M-L junctions in E13.5 epidermis. Mean with standard deviation are shown. Note similar recovery profiles at A-P and M-L junctions. n=18 traces each of Fz6-3XGFP and tdT-Vangl2 at M-L junctions. **(E and F)** Plots depicting the half-life of recovery of Fz6-3XGFP (E) and tdTomato-Vangl2 (F) with median and inter-quartile range shown. Pairwise comparisons are not significantly different except where noted. Half-life values were computed only from the subset of traces that fitted a one-phase association equation with a chi squared value of $\geq 0.8$. Scale bar: 5μm.

**Figure S4**

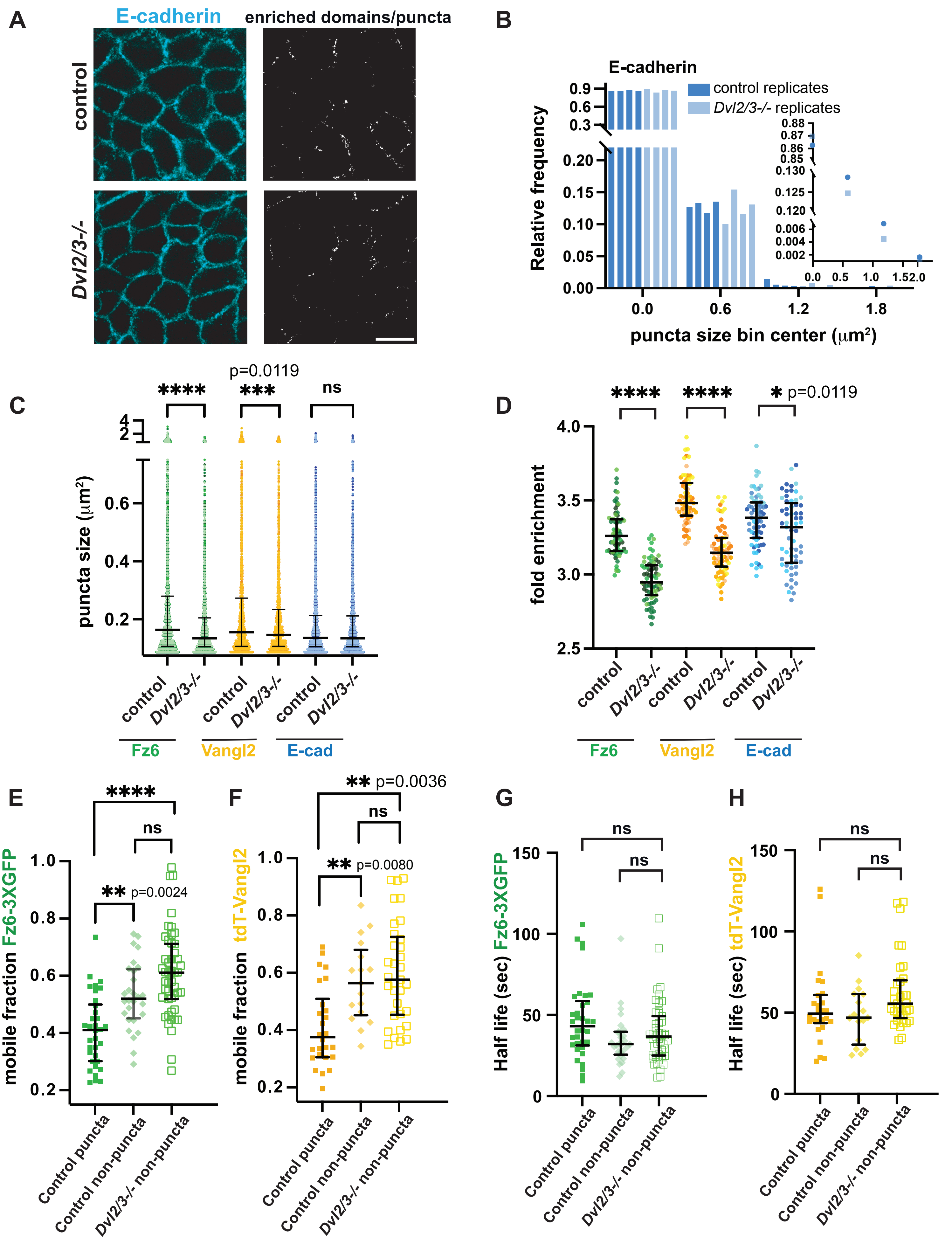
A
E-cadherin
enriched domains/puncta
control
Dvl2/3-/-
B
E-cadherin
control replicates
Dvl2/3-/- replicates
Relative frequency
puncta size bin center (μm²)
C
p=0.0119
****
***
ns
puncta size (μm²)
control
Dvl2/3-/-
Fz6
Vangl2
E-cad
D
****
****
* p=0.0119
fold enrichment
control
Dvl2/3-/-
Fz6
Vangl2
E-cad
E
****
ns
** p=0.0024
mobile fraction Fz6-3XGFP
Control puncta
Control non-puncta
Dvl2/3-/- non-puncta
F
** p=0.0036
ns
** p=0.0080
mobile fraction tdT-Vangl2
Control puncta
Control non-puncta
Dvl2/3-/- non-puncta
G
ns
ns
Half life (sec) Fz6-3XGFP
Control puncta
Control non-puncta
Dvl2/3-/- non-puncta
H
ns
ns
Half life (sec) tdT-Vangl2
Control puncta
Control non-puncta
Dvl2/3-/- non-puncta

**Supplementary Figure 4**. **Additional analysis of puncta formation by Fz6, Vangl2 and E-cadherin (A)** Planar views of epidermal basal layer of E15.5 embryonic skin immunostained for E-cadherin (left). Right panels show corresponding segmented images depicting spatially enriched domains/puncta of E-cadherin. **(B)** Normalized frequency distribution of E-cadherin puncta sizes. Each bar represents frequency of one control or *Dvl2/3-/-* replicate epidermis from 4 embryos. Inset shows relative frequency distribution of puncta sizes pooled from 4 control and 4 *Dvl2/3-/-* samples. p=0.6752 by Chi-Square test. n= 2549 puncta pooled from 4 control skins, n=2245 puncta pooled from 4 *Dvl2/3-/-* skins. **(C)** Plot showing spread of puncta sizes with median and interquartile range for Fz6, Vangl2 and E-Cadherin in control versus *Dvl2/3-/-* epidermis. **(D)** Fold enrichment of protein in puncta per ROI (ratio of mean intensity within puncta to mean intensity of the entire junctional network in the ROI) for Fz6, Vangl2 and E-cadherin in control versus *Dvl2/3-/-* epidermis. Median and interquartile range are shown. n= 69 ROIs of Fz6, Vangl2 and E-cadherin pooled from 4 control embryos and n= 71 ROIs of Fz6 and Vangl2 and 61 ROIs of E-cadherin pooled from 4 *Dvl2/3-/-* embryos. **** indicates p<0.0001 by KS test. **(E, F)** Mobile fractions of Fz6-3XGFP (E) and tdTomato-Vangl2 (F) from FRAP traces in Figure 4G and 4H. Mobile fractions were computed from the subset of traces that fitted to a one-phase association equation with a chi-squared value of ≥ 0.8. Median and interquartile range are shown. **(G, H)** Half-life of recovery of Fz6-3XGFP (G) and tdTomato-Vangl2 (H) from FRAP traces in Figure 4G and 4H. Half-lives were computed only from the subset of traces that fitted with a chi squared value of ≥ 0.8 to a one-phase association equation. **** indicates p<0.0001, tested by KS test in C-H.

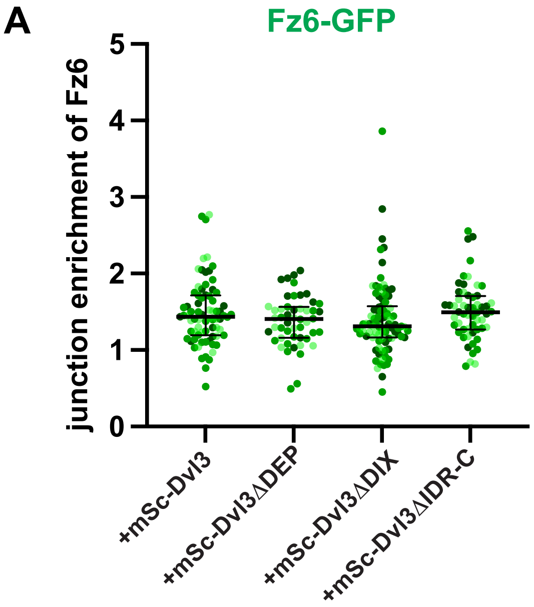
A
Fz6-GFP
junction enrichment of Fz6
5
4
3
2
1
0
+mSc-Dvl3
+mSc-Dvl3ΔDEP
+mSc-Dvl3ΔDIX
+mSc-Dvl3ΔIDR-C

**Supplementary Figure 5**. **Fz6 is accumulated to the same extent when co-expressed with full-length or domain-deleted Dvl3 mutants. (A)** Junction enrichment ratio of Fz6-GFP at keratinocyte junctions co-expressed with either full length or domain-deleted Dvl3 mutants. Median and interquartile range are shown. Note Fz6-3xGFP enrichment is similar across all conditions suggesting the lower recruitment of certain Dvl3 mutants is not due to lower levels of Fz6 at junctions. n= 75 junctions for Dvl3, 50 junctions for Dvl3ΔDEP, 88 junctions for Dvl3ΔDIX and 56 junctions of Dvl3ΔIDR-C, pooled from three replicates.

### Figure S6

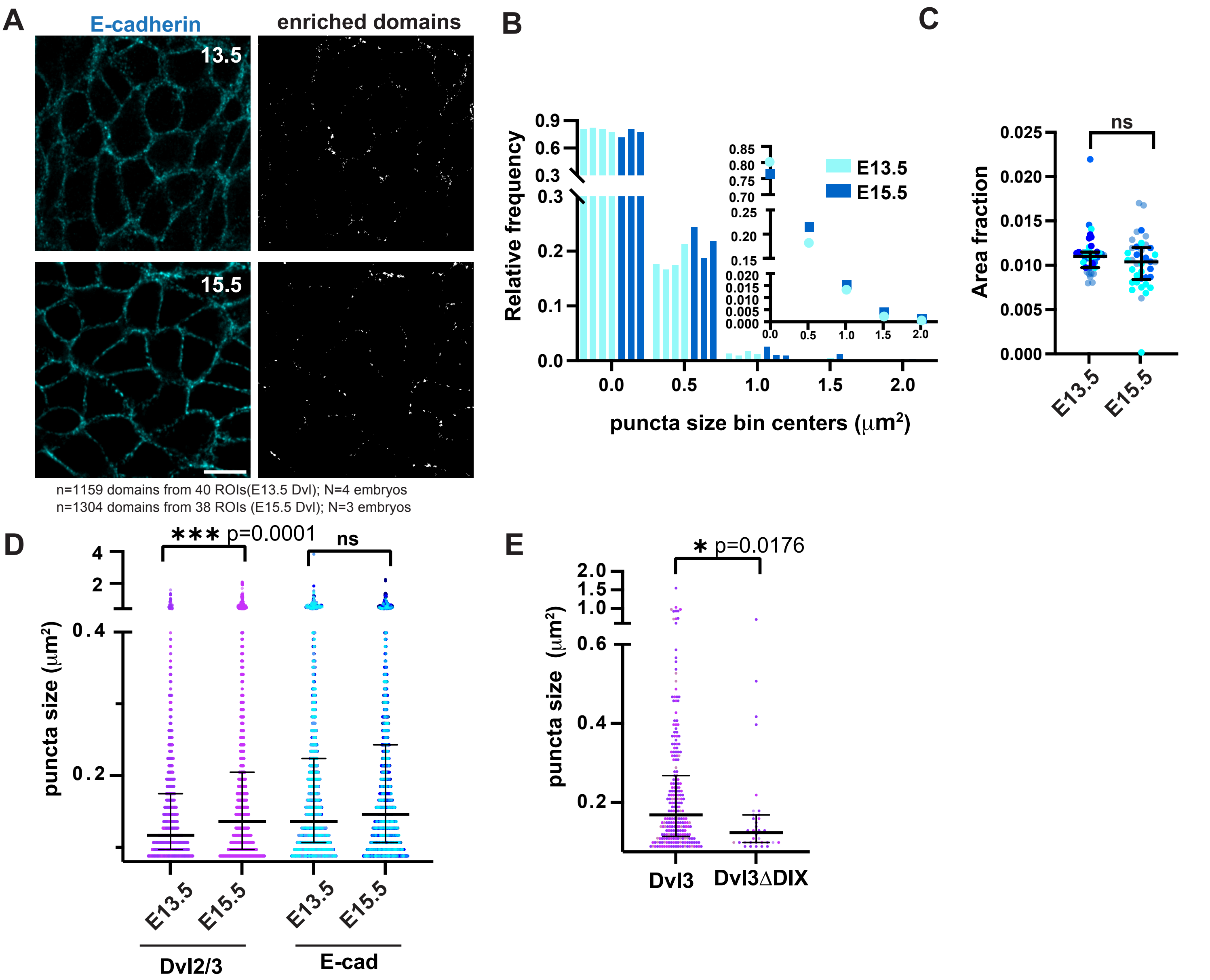
A
E-cadherin
enriched domains
13.5
15.5
n=1159 domains from 40 ROIs(E13.5 Dvl); N=4 embryos
n=1304 domains from 38 ROIs (E15.5 Dvl); N=3 embryos
B
Relative frequency
puncta size bin centers (µm²)
E13.5
E15.5
C
Area fraction
ns
E13.5
E15.5
D
puncta size (µm²)
*** p=0.0001
ns
E13.5
E15.5
E13.5
E15.5
Dvl2/3
E-cad
E
puncta size (µm²)
* p=0.0176
Dvl3
Dvl3ΔDIX

**Supplementary Figure 6. Additional analysis of puncta formation by Dvl2/3 and E-cadherin.** **(A)** Planar views of basal layer from E13.5 and E15.5 epidermis stained with anti-E-cadherin antibody. Right panels show segmented enriched domains. Note that E-cad accumulation into enriched domains/puncta is similar between the two stages. **(B)** Normalized frequency distribution of E-cadherin puncta/enriched domain sizes, with each bar representing one replicate epidermis at E13.5 (light blue) and E15.5 (dark blue) from 4 and 3 embryos, respectively. The inset shows normalized frequency distribution all puncta sizes pooled from 4 E13.5 and 3 E15.5 embryos. Distributions are not significantly different by Chi-Square test. n= 1808 puncta/enriched domains from 4 embryos at E13.5 and 1495 puncta/enriched domains from 3 embryos at E15.5. **(C)** Percent junctional area covered by puncta/enriched domains of E-cadherin, with median and inter-quartile range. n= 40 ROIs from 4 embryos at E13.5 and n=38 ROIs from 3 embryos at E15.5. **(D)** Quantification of Dvl2/3 and E-cadherin puncta sizes in E13.5 versus E15.5 epidermis with median and inter-quartile range. n= 1808 puncta/enriched domains of E-cadherin and 1159 puncta/enriched domains of Dvl2/3 from 4 embryos at E13.5; n=1495 puncta/enriched domains of E-cadherin and 1304 puncta/enriched domains of Dvl2/3 from 3 embryos at E15.5. **(E)** Quantification of mSc-Dvl3 and mSc-Dvl3ΔDIX puncta sizes in lentiviral transduced basal cells at E15.5. Median and inter-quartile range are shown. n= 245 puncta from 89 cell clusters and single cells and 3 embryos for mSc-Dvl3; n= 34 puncta from 93 cell clusters and single cells and 4 embryos of mSc-Dvl3ΔDIX.
